# PERREO: An integrated pipeline for repetitive elements analysis enables the repeatome expression profiling in cancer

**DOI:** 10.64898/2026.04.08.714730

**Authors:** Francisco Rodríguez-Martín, Mario Masero-León, Daniel Gómez-Cabello

## Abstract

Transcriptome-wide profiling of repetitive elements expression reveals transposable element-derived transcripts that are deregulated in diverse biological contexts including cancer. However, most RNA-seq pipelines are optimized for annotated genes and substantially undercount repeat RNA molecules, limiting their discovery and characterization. Here we present PERREO, a comprehensive, user-friendly pipeline for analyzing repetitive RNA elements from short- and long-read sequencing data. PERREO performs quality control, repeat-aware alignment and quantification, differential expression analysis, co-expression network analysis, and de novo transcript assembly with minimal computational expertise required. We validate PERREO across cell lines, tumor tissues and liquid biopsies, demonstrating superior sensitivity to repetitive RNA signatures compared with standard RNA-seq approaches. PERREO integrates predictive modelling to identify biological associations and generates publication-ready visualizations. By removing the bioinformatic barrier to repetitive RNA discovery, this pipeline enables broader investigation of the repeatome’s role in cellular biology and disease, yielding valuable results that, for specific analytical objectives, outperform certain existing tools and pipelines.

## MAIN

Characterizing the full transcriptome requires systematic detection of repetitive RNA elements (repRNAs), a task that has proved challenging with short-read sequencing. Recent advances in long-read technologies (PacBio, Oxford Nanopore) have enabled better coverage of repetitive genomic regions, as exemplified by the T2T human genome project^1^. However, Illumina short-reads remain the most widely used platform, and neither technology has yielded a comprehensive, standardized pipeline for repeats’ expression profiling. Repeat annotation, quantification and statistical analysis can be carried out by different specialized software that follow distinct strategies to deal with these elements, such as RepEnrich^2^, TEtranscripts^3^ and SQuIRE^4^. However, to the best of our knowledge, none of the currently available tools for repRNAs analysis is designed to seamlessly handle heterogeneous RNA-seq data types from different sequencing technologies, nor do they integrate downstream coexpression network analyses and supervised classification models specifically aimed at evaluating the diagnostic and prognostic potential of repRNAs.

Transposable elements (TEs) and other repetitive DNA sequences represent major sources of genetic variation, contributing to copy number and structural variants, as well as to insertions and deletions. Moreover, they can influence gene transcription and splicing^5^. Repetitive elements are generally transcriptionally silent in inactive regions of the genome, but transposon activation and epigenomic deregulation alter their expression in various pathological contexts, particularly cancer^6^. Emerging evidence demonstrates that repetitive RNA enrichment in circulating cell-free RNA is associated with disease state, suggesting these molecules are valuable biomarkers^6^. Notably, specific TE-derived RNAs have already been implemented in clinical and translational applications: HERV-E transcripts have been used as highly selective targets in clear cell renal cell carcinoma^7^, and LINE-1 ORF1/ORF2 proteins have been deployed as prognostic and early-detection biomarkers in breast^8,9^, colorectal^9,10^, hepatocellular^9,11^ and ovarian cancer^9,12^, thereby strengthening their potential not only as diagnostic and prognostic biomarkers, but also as markers of disease progression and treatment response. Yet, despite their potential, few bioinformatic tools specifically optimize repeatome detection, forcing researchers without specialized expertise to either lose critical information or require extensive custom analysis. Here we present PERREO, a comprehensive, user-friendly pipeline for detecting and quantifying repetitive RNA elements across diverse sample types and sequencing technologies.

## RESULTS

Long-read sequencing technologies provide crucial support for improving the detection of repetitive elements, as they enable the construction of more complete genome assemblies that fill many of the remaining gaps in the GRCh38 assembly (Fig. 1a). We are focused on the identification of repRNAs in several cancer models due to relevance of these elements in initiation and tumor progression caused by genomic instability in cancer cells (Fig. 1b). RNAs, and specifically these elements, with high stability and repeat in the genome, arise as an improved biomarker for their stability in biofluids, deregulation in cancer cells and innovative techniques to be detected with high sensitivity. PERREO not only identifies repRNAs but also integrates repeat-aware alignment, differential expression analysis, transcriptome assembly, coexpression network inference and predictive modeling, and is compatible with emerging reference genomes such as T2T-CHM13 (Fig. 1c shows the complete workflow). We demonstrate that this framework offers clear advantages over existing software-based approaches and pipelines in terms of flexibility, interpretability and analytical scope, and it also enables the discovery of novel repeat-containing transcripts and circulating repRNAs. Hence, PERREO provides a practical, reproducible entry point for researchers from diverse backgrounds to explore the contribution of repetitive sequences to cancer biology.

**Fig. 1:**
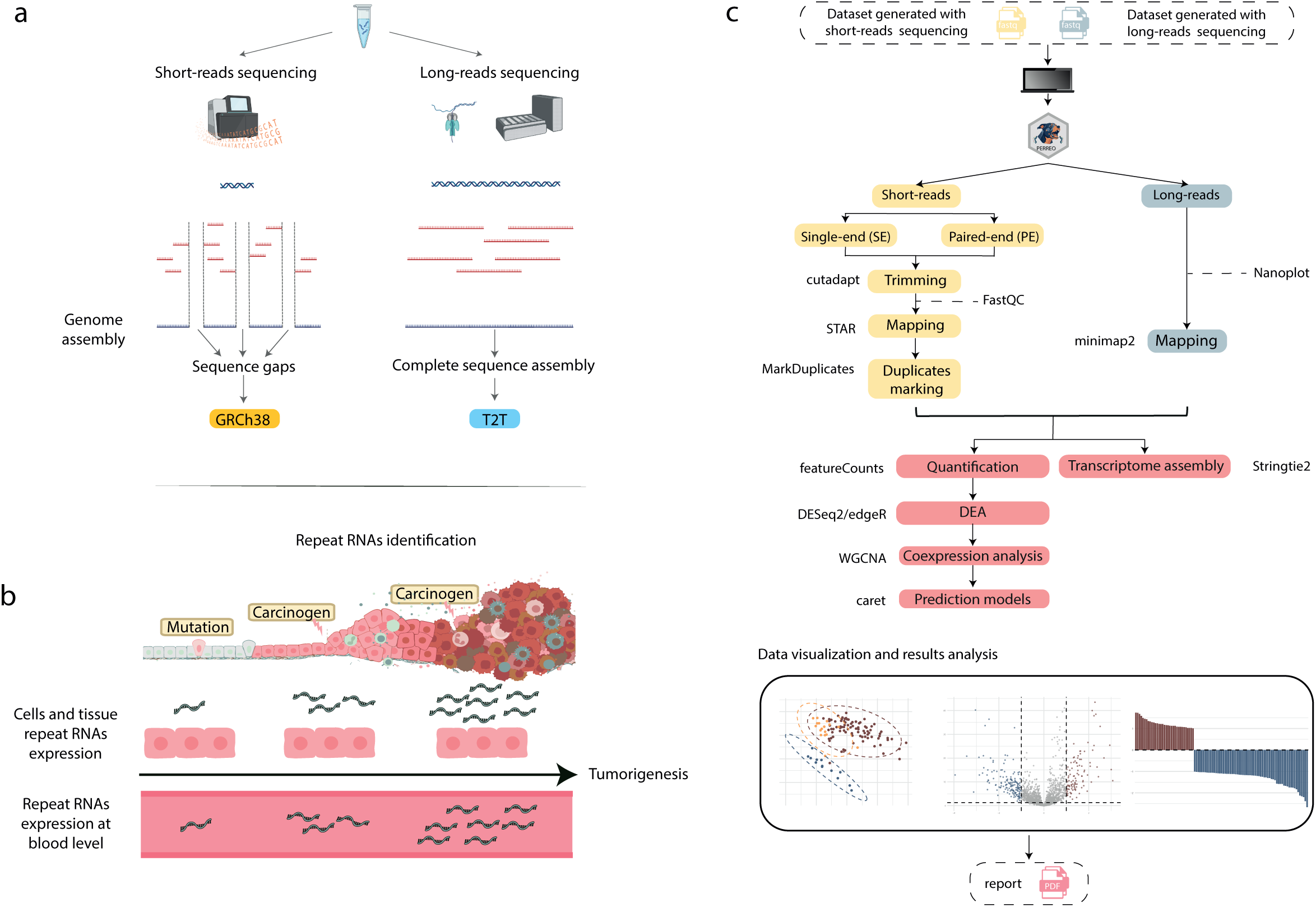
PERREO main pipelines and short-reads and long-reads sequencing technologies-based assemblies. **a,** GRCh38 and T2T-CHM13 genome assemblies built using short-reads and long-reads sequencing technologies, respectively. **b,** Specfic repRNAs expression increase as tumorigenesis and tumor progression takes place in cancer contexts because of genome instability. Parallelly, repRNAs show significant alterations in the cell-free transcriptome as these molecules can be released to blood. **c,** Summary of PERREO workflows. Different pipelines are implemented for data derived from long-reads experiments and paired-end and single-end sequencing generated with short-read approaches. Computational workflow generates a pdf report including different plots to interpret the results.

PERREO is designed as a modular, one-command-execution pipeline that automates the complete workflow previously described. The pipeline accepts raw sequencing data (FASTQ format), together with user-provided reference genome, genomic and repeat annotations, adapter sequences, and configuration files. PERREO generates three main analysis modes to accommodate different sequencing technologies: <u>SR-PE mode.</u> Paired-end short-read RNA-seq data analysis focused on repeat sequences, using the STAR^13^ aligner and a featureCounts^14^ configuration tailored to the handling of multi-mapping reads.

<u>SR-SE mode.</u> Single-end short-read RNA-seq data analysis applying an analogous processing strategy, very similar to SR-PE mode.

<u>LR mode.</u> Direct RNA-seq (Oxford Nanopore long-read) data analysis focused on long repeat-derived RNAs, using the minimap2 aligner^15^, Nanopore-specific processing tools and featureCounts with long-read–specific settings.

PERREO enables reporting of multi-mapping reads at the alignment step for both STAR and minimap2, and featureCounts consistently handles them by fractionally assigning counts across the genomic loci to which each read aligns, regardless of the analysis mode being run.

All modes output a standardized count matrix compatible with R-based differential expression analysis (edgeR^16^, DESeq2^17^) and generate automated quality reports and publication-ready visualizations. The pipeline is in process of being containerized in Bioconda for reproducibility and is organism-agnostic, requiring only appropriate reference files (Fig. 1c shows the complete workflow).

### Design and overview of PERREO

Comprehensive characterization of the repetitive transcriptome requires specialized handling of multi-mapping reads and repeat-specific genomic annotations and functionalities that standard RNA-seq pipelines systematically overlook or actively discard. We developed PERREO as a unified, modular, and containerized pipeline that processes raw sequencing data through three technology-specific pathways while converging at a shared downstream analysis framework, enabling seamless analysis of repetitive RNA elements across diverse sample types and sequencing modalities without requiring specialized bioinformatic expertise.

A critical design principle underlying PERREO is organism-agnosticity combined with reference flexibility. The pipeline embeds no hard-coded organism-specific parameters or annotation databases, instead accepting user-supplied references. This design choice confers two substantial advantages. First, it enables cross-species application, from humans to model organisms extensively used in cancer research (mice, zebrafish, Caenorhabditis elegans) to non-model organisms, using identical pipeline code. Second, PERREO is future-proof because it can adapt to ongoing improvements in genomic references. As shown by the recent completion of the T2T (Telomere-to-Telomere) human genome, more complete and accurate reference assemblies continue to emerge over time. Rather than requiring pipeline updates, PERREO users simply provide updated reference and annotation files, enabling immediate leverage of improved genomic resources.

The pipeline produces a set of unified, publication-ready outputs. It generates a comma-separated count matrix with both raw and normalized feature counts across all samples, which can be easily imported into statistical software or used in downstream analyses. For each sample, BAM alignment files are also provided, preserving full alignment information for more customized analyses. An automated quality control report, generated via MultiQC^18^ and FastQC^19^, provides summary statistics on sequencing quality, mapping efficiency, and repeat coverage. Differential expression results are exported as delimited tables containing log_2_ fold-changes (log_2_FC), adjusted p-values (FDR), and feature counts. The pipeline also generates a range of visualizations such as volcano plots, hierarchical heatmaps, principal component analysis plots, co-expression networks, and machine learning feature importance rankings, all generated in publication-ready formats. Finally, detailed execution logs and parameter records are included to ensure full reproducibility and clear documentation of the methodology.

By consolidating repeat-aware alignment, multi-platform compatibility, streamlined statistical analysis, and machine learning-based discovery into a single, containerized, user-friendly package, PERREO substantially lowers the technical barrier to repeatome research. This enables research groups lacking specialized bioinformatic expertise in repetitive element analysis to leverage the emerging clinical and biological significance of the repeatome, multiplying the pace of discovery in cancer biology, circulating biomarker development, and fundamental transcriptome characterization. To stablish the rationale for our repeatome discovery and characterization by PERREO in representative RNA-seq data samples from several sequencing platform and biological material. PERREO can analyse the repeat transcriptome profile associated with different cancer types derived from distinct types of samples, like plasma, tissue, and cell lines. To test the pipeline performance, we carried out bioinformatic analysis of publicly available data stored in different repositories, analyzing 330 samples in all. We downloaded five data sets of short-reads RNA sequencing experiments related to different types of samples and diseases in mice, dogs and humans. In addition, we also analysed long-read sequencing data from 4 cancer cell lines related to liver, leukocyte, breast, and colorectal cancer, and H9 human embryonic stem cell line (Fig. 2).

**Fig. 2:**
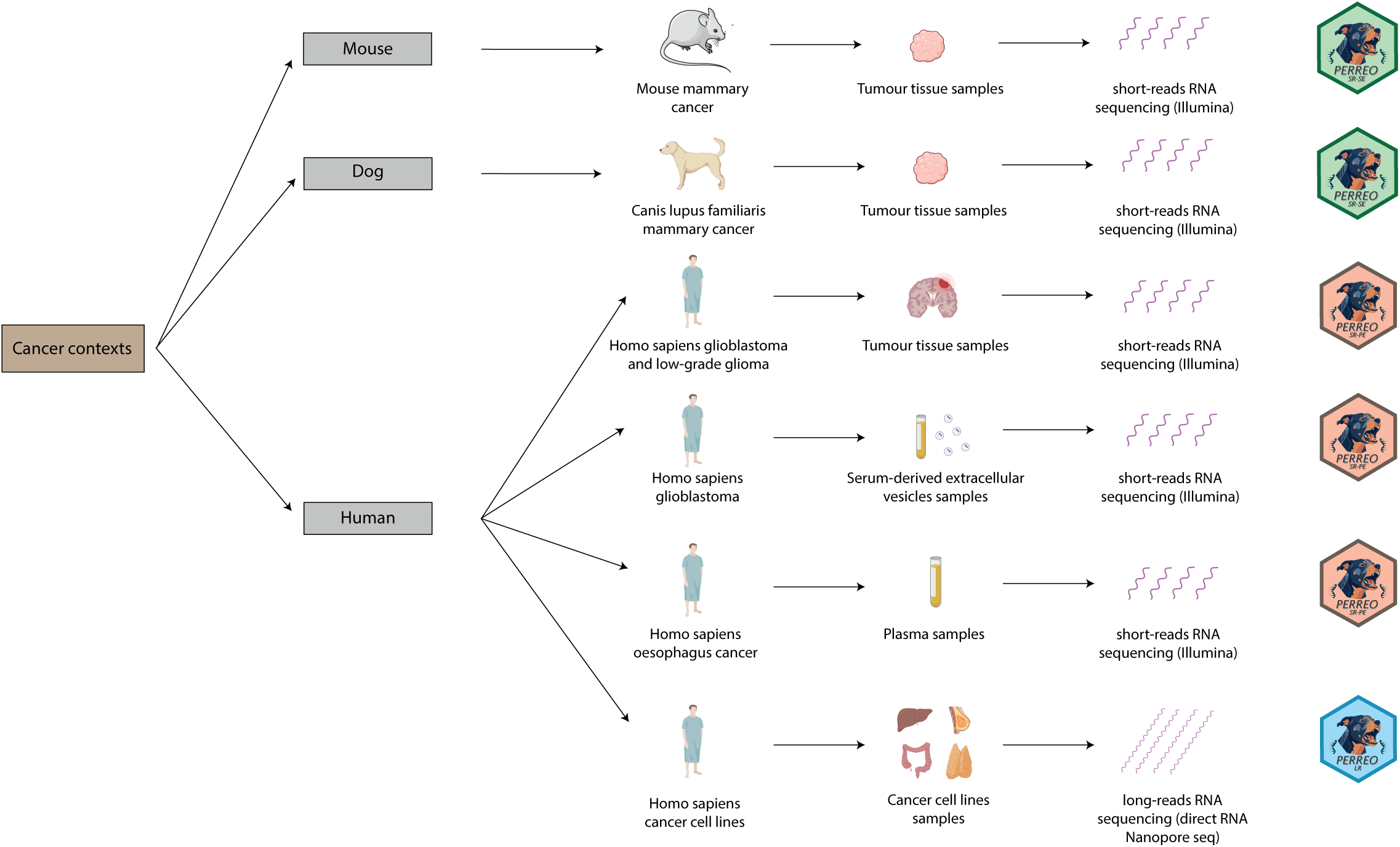
A Schematic summary of the workflows followed to analyze the repeatome expression profile from publicly available datasets. Datasets based on mice mammary cancer (N=16), dog mammary cancer (N=21), human glioblastoma and low-grade glioma tissues (N=115), serum-derived extracellular vesicles from human glioblastoma (N=111), plasma from oesophagus cancer patients (N=45), and different cancer cell lines (N=22).

### Plasma repRNAs as candidate biomarkers in oesophageal cancer

Cell-free RNA circulating in plasma represents a minimally invasive source for disease biomarker discovery, yet its fragmented nature and low abundance pose distinct analytical challenges^20^. We applied PERREO to systematically profile the repeat-derived transcriptome in plasma from oesophageal cancer (ESCA) patients and healthy controls (HC), processing data from 45 individuals (23 ESCA, 22 HC) stored in Gene Expression Omnibus (GEO) under GSE174302 accession code, using the short-read paired-end mode with default parameters (|log_2_FC| > 1, FDR < 0.05).

Mean repRNAs expression showed no statistically significant differences between ESCA and HC samples (Wilcoxon test, p > 0.05), consistent with expectations that repeat transcripts may constitute a relatively stable component of circulating RNA (Fig. 3a). Initial analysis revealed 448 repeat elements detected across all samples, with distribution dominated by LINE/L1 elements (24.8% of detected features), followed by simple repeats (12.1%) and LTR retrotransposons (9.6%) (Fig. 3b). Differential expression analysis identified nine differentially expressed repeats (DERs), comprising six upregulated and three downregulated repRNAs in ESCA relative to HC samples (Fig. 3c,d). This signal demonstrates that despite overall repeatome stability at the categorical level, specific repeat elements respond to cancerous disease states.

**Fig. 3:**
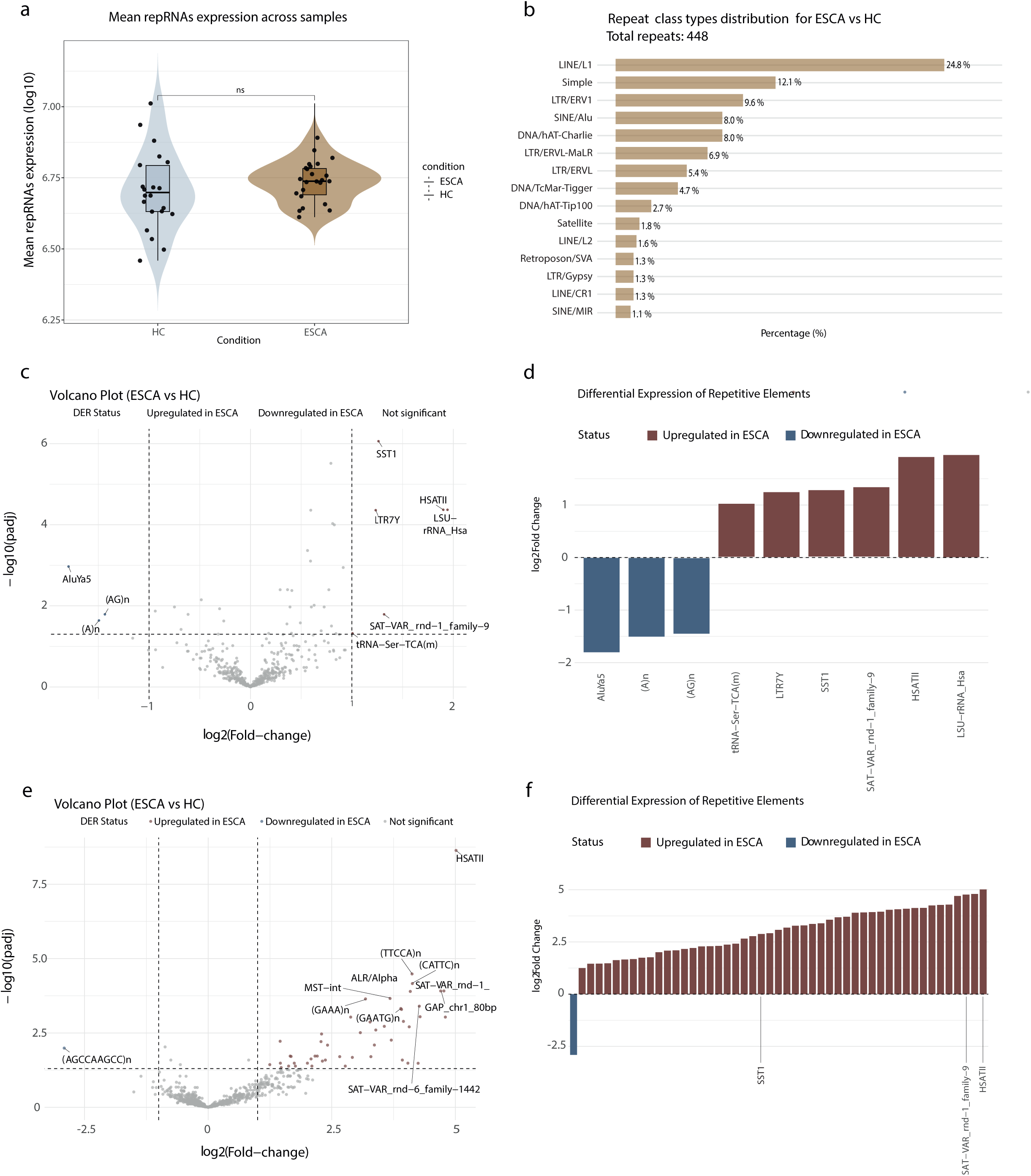
Oesophagous cancer cell-free transcriptome showed some repRNAs as biomarker candidates. **a,** Violin plot representing the median, the upper and lower quartiles of repRNAs mean expression in ESCA (N=23) and HC (N=22). Significance level was calculated with Wilcoxon test, ****P < 0.0001, ***P < 0.001, **P < 0.01, *P < 0.05, ‘ns’ P > 0.05. **b,** Most represented categories among the 448 features detected across study samples. **c, d,** Volcano plot where upregulated features in ESCA are colored in red and downregulated in ESCA are colored in blue **(c)** and Fold-change bar plots representing DERs **(d)** when procedence hospital was not included in the analysis. **e, f,** Volcano plot **(e)** and Fold-change bar plots representing DERs **(f)** when procedence hospital was included in the analysis.

We observed that hospital-of-origin metadata introduced batch effects that substantially modified results. When we re-analyzed a subset of previously analyzed samples with complete metadata and included hospital as a batch covariate in the statistical model, the differential expression signature strengthened dramatically: 48 upregulated repRNAs and only one downregulated repeat in ESCA versus HC (Fig. 3e, f). This substantial increase in sensitivity upon batch correction underscores the critical importance of PERREO’s automated batch effect handling, which would have been overlooked by researchers unaware of this technical covariate’s influence. The ability to specify batch variables through a simple metadata column represents a key practical advantage for users analyzing multi-site clinical cohorts.

### RepRNA signatures in glioblastoma tissue and extracellular vesicles

#### Impact of GRCh38 versus T2T-CHM13 on tissue repRNA profiles

The recent completion of the T2T (Telomere-to-Telomere) human genome assembly provides unprecedented representation of previously inaccessible repetitive regions. Next, we wanted to evaluate whether improved reference annotations materially impact repRNAs characterization using PERREO. We analyzed short-read RNA-seq data from glioblastoma (GBM, n=85), low-grade glioma (LGG, n=17), and healthy control brain tissues (HC, n=13) from the dataset stored under accession code GSE147352, by aligning reads against both the widely-used GRCh38 and the T2T-CHM13 assembly using identical PERREO parameters (|log_2_FC| > 1, FDR < 0.01).

Alignment quality metrics revealed substantial differences between reference genomes. T2T-CHM13-aligned samples contained 6.28% multimapped reads on average, compared with 11.30% for GRCh38-aligned data (Fig. 4a,b). This 45% reduction in multimapping ambiguity reflects T2T-CHM13’s superior ability to uniquely assign repetitive sequences to their correct genomic loci, a direct consequence of improved assembly of previously collapsed repetitive regions. This quantitative demonstration of reference quality impact validates a core design principle of PERREO: organism-agnosticity enabling users to immediately leverage improved genomic resources without pipeline re-implementation.

**Fig. 4:**
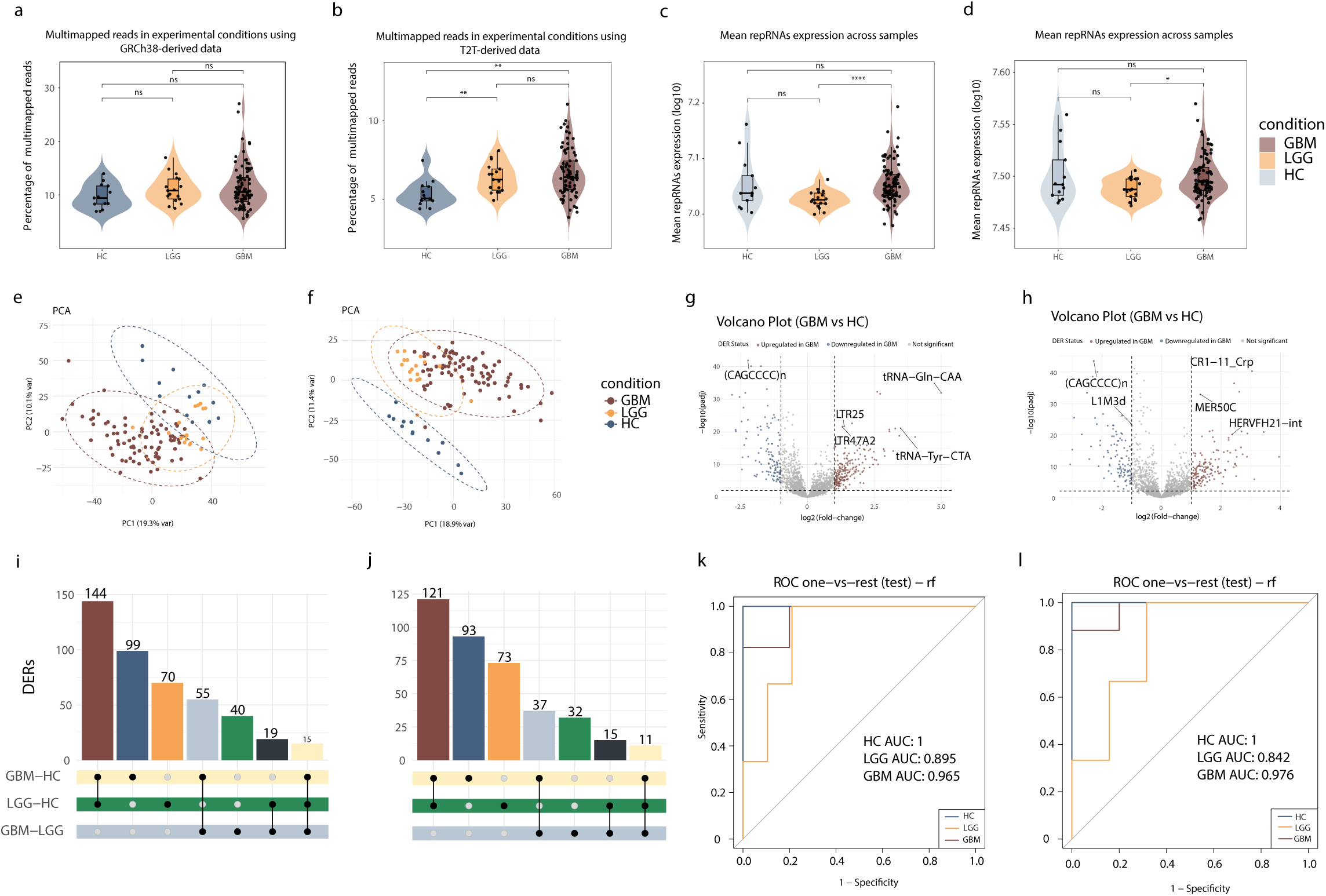
Results based on GRCh38 and T2T-CHM13 alignment show light variations in repRNAs differential expression search. **a, b,** Violin plots displaying the median, the upper and lower quartiles regarding the percentage of multimapped reads in HC (N=13), LGG (N=17) and GBM (N=85) for GRCh38-derived results **(a)** and T2T-CHM13 -derived results **(b)**. Significance level was calculated with Wilcoxon test, ****P < 0.0001, ***P < 0.001, **P < 0.01, *P < 0.05, ‘ns’ P > 0.05. **c, d,** Violin plots representing median, the upper and lower quartiles of mean repRNAs expression in each experimental condition using GRCh38 **(c)** and T2T-CHM13 **(d)** approaches. Statistical analysis was also realised with the Wilcoxon test. **e, f,** PCA plots obtained from GBM, LGG, and HC repRNAs using GRCh38 **(e)** and T2T-CHM13 **(f)** references. **g, h,** Volcano plots of GBM versus HC contrast using GRCh38 **(g)** and T2T-CHM13 **(h)** genomes, where the upregulated repRNAs in GBM are shown in red and the downregulated repRNAs are shown in blue. **i,j,** Upset plots showing the shared and unique DERs among the contrasts with GRCh38 **(i)** and T2T-CHM13 **(j)** references. **k, l,** ROC curves showing Random Forest algorithm performance on GRCh38-aligned-derived data **(k)** and T2T-CHM13-aligned-derived data **(l)**.

Regarding the mean repRNAs expression, we detected a significant increase in GBM with respect to LGG condition aligning reads against both GRCh38 and T2T-CHM13 assemblies (Fig. 4c,d). Interestingly, we also observed a light and non-significant decrease in repeats’ expression in LGG in contrast to HC, suggesting a minor reduction in repRNAs mean expression in less aggresive stages and a consequent rise in more aggresive states (Fig. 4c,d). Principal component analysis based on repeat element expression provided clearer separation between disease categories when using T2T-CHM13 references compared with GRCh38, with GBM and LGG samples forming more discrete clusters (Fig. 4e,f). These findings collectively demonstrate that PERREO’s flexibility to accommodate improved reference genomes has direct biological impact, and that advances in genome assembly quality translate to enhanced statistical power and biological clarity in repeatome studies.

Differential expression analysis (DESeq2, FDR < 0.01, |log_2_FC| > 1) revealed disease severity-associated variations in repeat element expression (Fig. 4g,h). Using T2T-CHM13 references, we detected 262 DERs in GBM versus HC comparison, 220 in LGG versus HC, and 95 in GBM versus LGG. Equivalent analyses with GRCh38 yielded higher DER counts (313, 248, and 129 respectively), reflecting the reference’s greater propensity to collapse repetitive sequences and potentially assign them spuriously. By intersecting the results from all contrasts, we found that many DERs were shared when comparing GBM and LGG against HC samples (Fig. 4i,j), suggesting that activation of some repeats may already occur at early stages of tumorigenesis. Specifically, 144 DERs and 121 DERs were common using GRCh38 and T2T-CHM13 assemblies, respectively. When we directly compared results between references, substantial overlap existed but with significant reference-specific components: 154 common DERs in GBM-HC comparisons (59% of T2T-CHM13 -derived DERs), 136 in LGG-HC (62%), and 52 in GBM-LGG (55%) (Extended Data Fig. 3). This pattern indicates that approximately 40-45% of detected differential repRNAs are reference-dependent, underscoring that biological conclusions drawn from repeatome expression analysis should account for reference choice.

PERREO supports predictive modeling using Random Forest and GLMnet classification algorithms, which demonstrated strong performance in terms of both accuracy and area under the ROC curve (AUC). For the GRCh38-based analysis, the Random Forest model achieved an accuracy of 0.909 and a multiclass AUC of 0.971. When using the T2T-CHM13 reference, accuracy remained the same (0.909) with a comparable AUC of 0.964 (Extended Data Fig. 4). To identify key contributors to model performance, we ranked the top 20 most important features from each Random Forest model and found that 8 of the 20 repRNAs were shared between the two references (Supplementary Table 10). Both models showed high performance, but they proposed novel repRNAs that did not overlap between the two models.

#### Extracellular vesicles-associated repRNAs in glioblastoma

Liquid biopsies from cancer patients represent one of the most promising approaches for early diagnosis and prognosis. They may reveal which upregulated repRNAs are released into the bloodstream via extracellular vesicles (EVs), potentially contributing to systemic communication in cancer. To explore this, we analyzed the dataset linked to GSE228512 code, using a slightly lower |log₂FC| threshold (> 0.8), accounting for the expected low RNA content in exosomes. A decrease in the mean expression of EV-derived repRNAs was observed in GBM samples compared with HC (Fig. 5a). Twenty DERs were identified, of which 18 were downregulated in GBM (Fig. 5b,c). Using this dataset, we developed classification models to distinguish between conditions. Here, the GLMnet algorithm outperformed Random Forest, achieving an AUC of 0.80 versus 0.75, respectively (Fig. 5d). Accordingly, the repeat-based predictors proposed by the two algorithms did not overlap. Interestingly, when comparing Random Forest models built from tissue and serum-derived EV data in GBM, one repeat feature appeared among the top 20 most important variables in both cases, which was the simple repeat (TGTTTT)n (Supplementary Table 10).

**Fig. 5:**
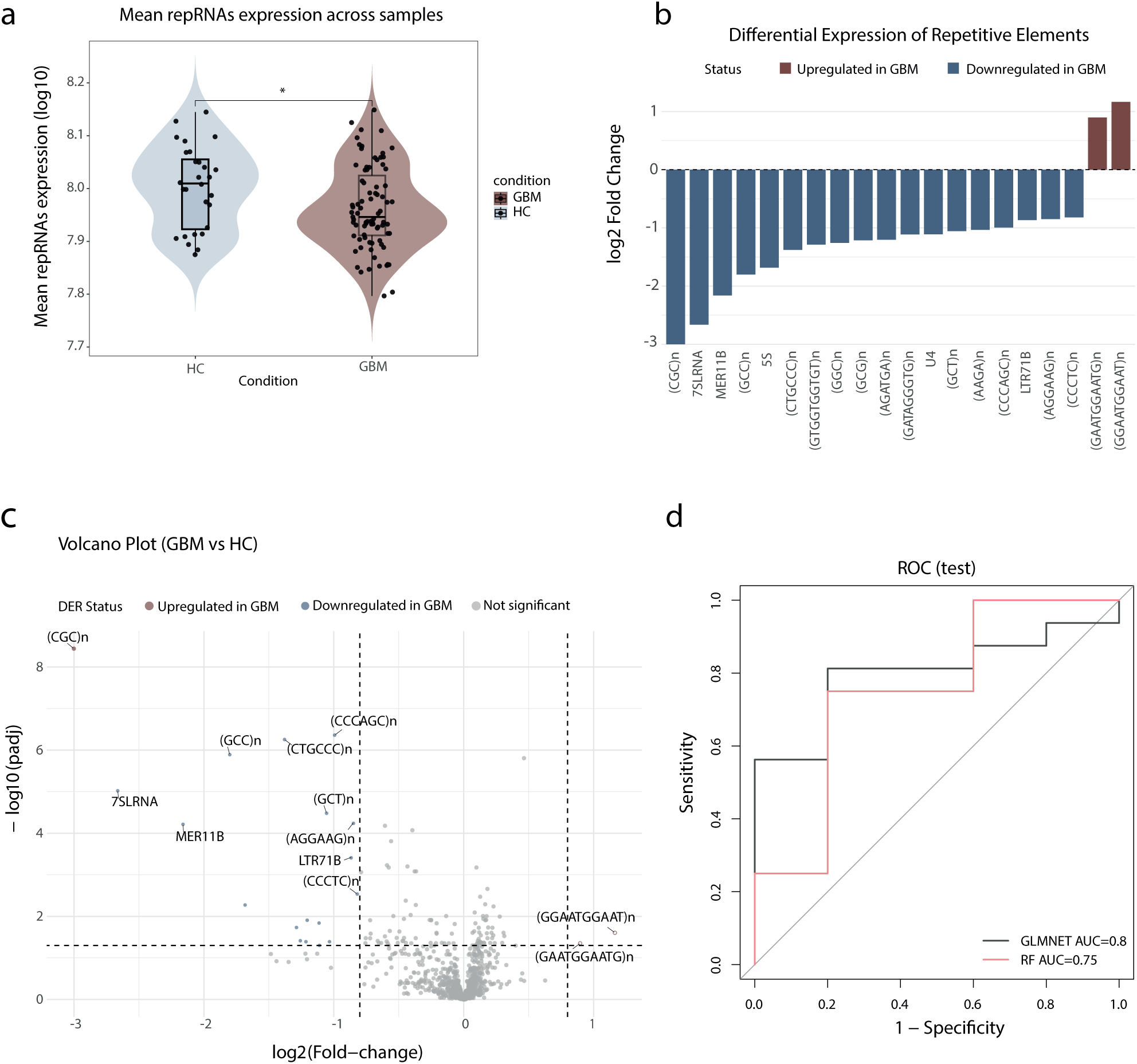
RepRNAs are detectable at extracellular vesicles obtained from plasma and increase their potential as biomarkers. **a,** Violin plot showing the median, the upper and lower quartiles of repRNAs mean expression in extracellular vesicles of GBM and HC samples. Significance level was calculated with Wilcoxon test, ****P < 0.0001, ***P < 0.001, **P < 0.01, *P < 0.05, ‘ns’ P > 0.05. **b,** Log2Fold-Change bar plot of DERs in GBM versus HC contrast. **c,** Volcano plot of GBM versus HC contrast, highlighting in red the upregulated repRNAs in GBM, and in blue the downregulated repRNAs. **d,** ROC curve of Random Forest and GLMnet models based on the performance (AUC).

### Long-read RNA sequencing reveals specific repeat transcriptome patterns across cancer cell lines

In view of the increasing importance of long-read sequencing technologies to detect centromeric and telomeric regions, we applied PERREO to long-read data to showcase the potential of the pipeline. We analyzed direct RNA sequencing data obtained from the Singapore Nanopore Expression project^21^, generated with Oxford Nanopore technology from several cancer cell lines, colon (HCT116), liver (HepG2), breast (MCF7), and leukemia (K562), using our pipeline configured for long-read data analysis (–batch no, –method edgeR, and default differential expression parameters).

In terms of repRNAs distribution, expression patterns varied markedly among the cell lines. Wilcoxon tests revealed significant differences in the contrasts K562 versus H9, K562 versus HepG2, MCF7 versus K562, and MCF7 versus HepG2. Among all, K562 exhibited the highest mean expression of identified repRNAs (Fig. 6a). The immune-related nature of K562 cells may underlie their distinct repeatome expression profile, as the other cell types analyzed are carcinoma-derived cell lines.

**Fig. 6:**
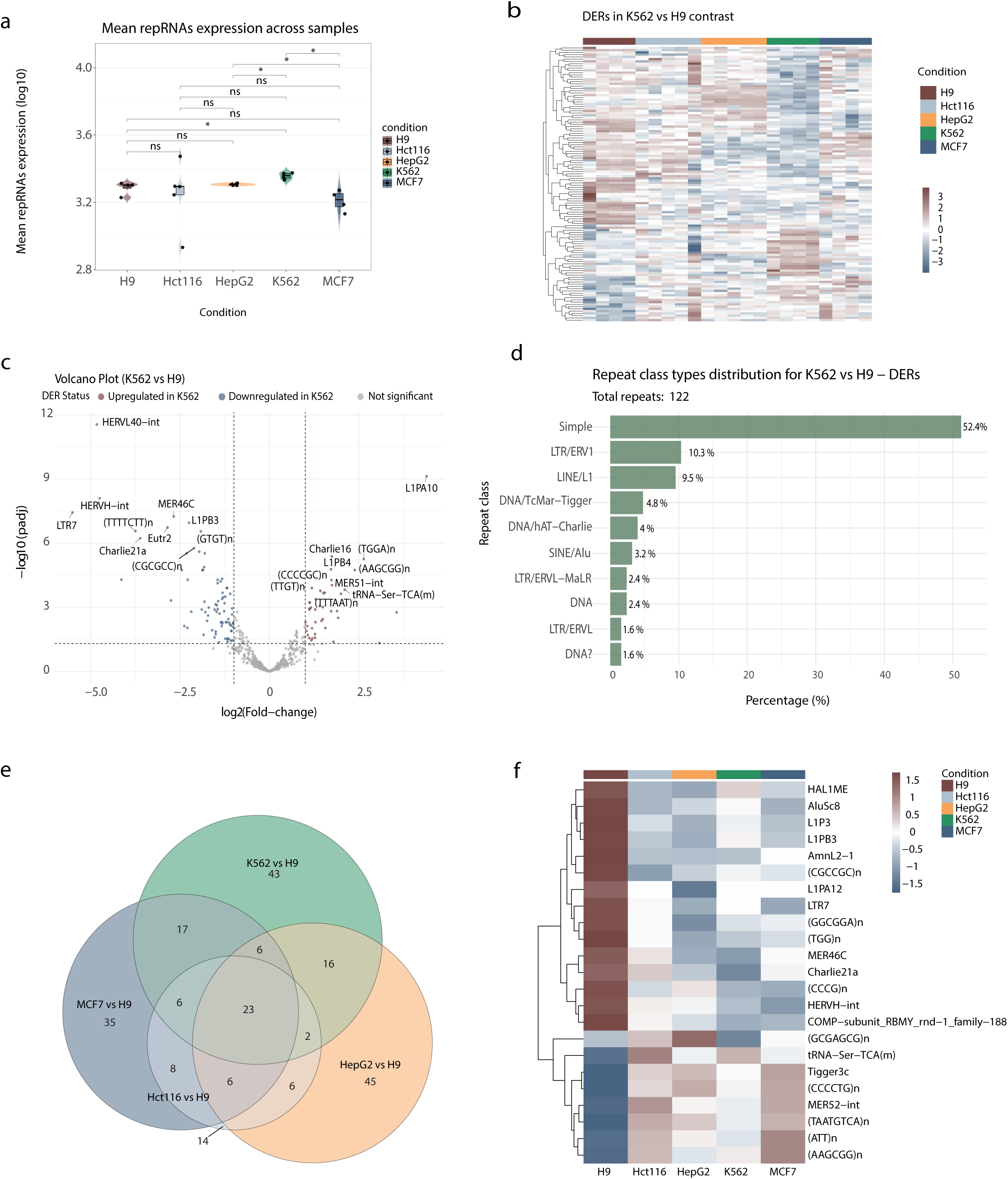
DirectRNA Nanopore long-sequencing technology shows that repRNAs expression profiles vary in different cancer cell lines. **a,** Violin plot showing the median, upper and lower quartiles of repRNAs expression in the five cell lines studied. Significance level was calculated with Wilcoxon test, ****P < 0.0001, ***P < 0.001, **P < 0.01, *P < 0.05, ‘ns’ P > 0.05. **b,** K562 versus H9 DERs expression heatmap. **c,** Volcano plot highlighting K562 versus H9 DERs. Upregulated repRNAs in K562 are shown in red, while downregulated repRNAs are shown in blue. **d,** Most represented repeat classes in 122 DERs in K562 versus H9 contrast. **e,** Euler diagram of all contrasts where cancer cell lines are compared with H9 line. **f,** Mean expression heatmap of the 23 DERs identified in all the cancer cell lines versus H9 line.

Interestingly, when statistically comparing all cell lines, the most divergent repeatome expression profiles corresponded to H9 and K562, with 122 DERs showing a consistent expression pattern across samples and predominantly upregulated in H9 relative to K562 (Fig. 6b,c). Some of these repRNAs belong to transposable element subfamilies associated with genome instability and cancer, such as L1PA10 and L1PB4, which are part a lineage of LINE-1 elements. These retroposons are usually aberrantly activated in cancer contexts and are potential diagnostic markers and therapeutic targets^22^. More than half of DERs identified in this contrast corresponded to simple repeats category, constituting the most represented class, followed by LTRs and LINE/L1 family (Fig. 6d). Most DERs identified in the rest of the contrasts of this study and in glioblastoma at tissue and EVs levels were also associated with simple repeats class, suggesting this group as the most altered repeat type in cancer contexts.

Importantly, beyond cell line–specific differences, we identified 23 repeat features that were consistently differentially expressed across all cancer cell lines compared with the non-cancerous H9 line (Fig. 6e,f). This consistency suggests a shared repeatome signature associated with cancer phenotypes that could be mediated by some of these DERs. Notably, a substantial fraction of the repRNAs upregulated in cancer cell lines corresponded to simple repeats. In contrast, specific elements from the LINE-1 lineage were upregulated in H9, which is consistent with the idea that LINE-1 elements are more active in early developmental contexts characterized by hypomethylation of these genomic regions^23^.

## Discussion

The potential of repRNAs as diagnosis, prognosis and stratification tools has been largely underestimated, as traditional RNA-seq analyses usually ignore repetitive sequences due to their historical classification as “non-functional” and their predominant localization in non-coding regions. In addition, these sequences are frequently discarded as noise when researchers focus on protein-coding transcripts, since their high multimapping rate is typically considered a technical challenge. We addressed this gap generating PERREO, a user-friendly pipeline to analyse RNA-seq data to determine these repRNAs on different biological models. Considering PERREO-derived results, repRNAs can classify, stratifying and predicting phenoypes at tissue, blood, EVs and cell lines level, similarly to genome coding sequences.

Long-read sequencing technologies have greatly expanded our ability to detect repetitive elements, owing to their high read length^24^. However, several practical challenges remain, as current protocols are not yet fully optimized for repeat quantification. In the case of Oxford Nanopore direct RNA sequencing, most standard approaches require reads to be polyadenylated, which inherently biases detection against non-polyadenylated repRNAs ^25^. Recently, alternative methods have emerged to capture RNAs regardless of their polyadenylation status, such as Nano3P-seq, which uses template-switching technology and avoids poly(A) selection^26^.

Against this backdrop, our work advocates for automated, publicly available pipelines to explore repRNAs expression dynamics by leveraging the vast number of existing short-read RNA-seq datasets together with already published long-read data. The growing diversity of high-quality reference genomes and curated annotations further enables complementary strategies to identify repRNAs and to perform downstream analyses in a more robust and reproducible manner.

We propose that combining differential expression analysis, coexpression analysis, transcriptome assembly and predictive modeling can substantially enhance the utility of repRNAs profiling in oncology. This integrated framework not only improves our ability to address complex biological questions but also lowers the technical barrier for researchers without extensive programming expertise, thereby increasing the likelihood of identifying repeat-based biomarkers for early cancer diagnosis. At the same time, transcriptome assembly approaches expand the discovery of novel transcripts and provide insight into the biological roles of repRNAs and the regulatory context that drives their transcription.

We benchmarked PERREO against other related software tools (Supplementary Note 4). Firstly, we attempted to compare STAR pipeline proposed in PERREO against HISAT2^27^ mapper and Salmon pseudoaligner^28^. However, when mapping with HISAT2 multiple samples failed at the alignment step using identical resources, maybe as consequence of the high multimapping burden and the specific indicated values for some parameters, leading us to conclude that STAR performance overpasses HISAT2 capacity to deal with multimappers. On the other hand, Salmon showed faster alignment using the same resources, but in this case, due to its transcriptome-focused approach, genomic coordinates information was lost, what limited our interpretation of genomic origin of these transcripts. The Salmon developers do not recommend removing duplicates when unique molecular identifiers (UMIs) are not included in the experimental design, so we retained duplicates when running Salmon. In parallel, we executed PERREO under two settings, either retaining or removing duplicates. However, the results obtained with PERREO using the remove_duplicates *yes option* were more like those produced by Salmon. PERREO found 49 DERs, while Salmon identified 47, from which 10 were statistically significant in both pipelines (Extended Data Fig. 5b). Salmon redistributes multi-mapping reads and models several technical biases, yielding abundance estimates that tend to down-weight redundant signal in a way similar to explicit PCR duplicate removal. This behavior likely underlies the concordance we observe between the Salmon pipeline without duplicate removal and the PERREO workflow in which PCR duplicates are explicitly removed.

Then, we also compared PERREO performance with TEtranscripts, a tool specifically designed for Transposable Elements (TE) analysis, using T2T-CHM13 pre-built annotations^3^. In PERREO, multi-mapping reads are included and distributed fractionally across RepeatMasker^29^ gene_ids using featureCounts (1/n per alignment), whereas TEtranscripts further refines multi-mapper assignment with specific algorithm know as Expectation-Maximization (EM)^30^ to infer the active copy at the TE subfamily level. In our dataset, PERREO produced a larger count matrix (2,393 features after stringent filtering) compared with TEtranscripts (1,283 features) and identified 262 DERs versus 47 detected by TEtranscripts, with 11 shared between both approaches (Extended Data Fig. 6b). This difference in multi-mapping handling and repeat annotations likely explains a substantial part of the discrepancies we observe between both repeat quantification strategies. Additionally, PERREO exhibited substantially shorter runtimes (less than 4 hours, compared with almost 27 hours for TEtranscripts), consistent with its lower algorithmic complexity and the absence of iterative EM-based optimization steps. This constitutes an important point as PERREO is faster and less resource-intensive (Extended Data Fig. 6c & Supplementary Note 4.2).

Handling of multi-mapping reads in RNA-seq has been extensively reviewed, and equal fractional assignment is explicitly described as a standard strategy for aggregated features such as genes or repeat families, alongside more complex rescue and Expectation-Maximization-based approaches^31^. Because our primary endpoint is repeat expression at the RepeatMasker gene_id (subfamily) level rather than locus-specific activity, we deliberately adopt this simple 1/n fractional assignment, which is transparent, easy to reproduce, fast and conceptually aligned with these recommendations. In parallel, TEtranscripts is fully compatible with this pipeline and could provide a complementary approach to identify specific transcriptionally active copies.

Within this study, we observed that repeat expression profiles are altered in cancer, although changes in mean expression of repetitive elements are very variable in most cases. Some of these changes were detectable in blood-derived samples, which reinforces their potential as candidate biomarkers, even though their functional contribution in these contexts remains poorly understood. In GBM patient samples, we found that repRNAs expression varied not only at the tissue level but that specific repRNAs could also be detected within extracellular vesicles. Interestingly, the correspondence between these two different cohorts analyzed by PERREO identify one candidate repRNA ranked among the most important variables in models derived from both tissue and data. This may be explained by glioma tumor heterogeneity and the presence of the blood–brain barrier (BBB), which together complicate the discovery of biomarkers for disease diagnosis and prognosis and make this task highly challenging^32^. From a broader perspective, we also identified repRNAs that appeared consistently altered across multiple cell line datasets, suggesting the existence of shared repeat-associated signatures, as were found between different cancer cell lines (Fig. 6f). In cancer, the transcriptional activity of specific repetitive and transposable elements, such as the SST1 macrosatellite, has clinical relevance because deregulated expression of these elements can directly compromise genomic integrity and stability^5^. Pericentromeric HSATII upregulation has been described as a common feature of epithelial cell–derived cancers^33^, and we also detected its overexpression in plasma from ESCA patients as well as we also identified SST1 upregulation in these patients. Although GBM and LGG are not of epithelial origin, both tumor types showed increased HSATII levels compared with healthy controls. Furthermore, several HERV subfamilies have been reported as deregulated in glioblastoma tissue, including both upregulated and downregulated^34^, and we observed significant upregulation of HERVK13-int, HERVFH21-int, HERV3-int, HERV4_I and HERVI-int, together with downregulation of HERV9N-int in GBM samples relative to healthy brain tissue.

Despite the remaining uncertainty, previous work has also shown that repRNAs can promote mesenchymal gene expression programs in pancreatic ductal adenocarcinoma cells (PDAC) and influence tumor microenvironment cell-state transitions and crosstalk via extracellular vesicle release, underscoring the importance of investigating their role in cancer dynamics, beyond their potential as biomarkers^35^.

In conclusion, we developed PERREO for systematic repRNA discovery and repeatome expression profiling, providing a framework to identify novel repeat-derived biomarkers and to facilitate the broader adoption of repeat-aware transcriptomic analyses within the research community. The fact that PERREO is applicable to multiple species further increases its potential, as it can be used to interrogate oncological contexts in widely employed model organisms such as yeast and mouse. Beyond oncology, it is likely that other disease settings characterized by genome instability also involve altered expression of repetitive elements as it occurs in neurodegenerative diseases^34^. In such scenarios, repRNAs may act not only as passive and early markers of dregulation, but as active components shaping transcriptome dynamics in disease with huge prognostic and diagnostic potential.

## Methods

### Pre-analysis requirements

The input sample sheet must include, for each sample, a unique identifier, the library strandedness, and the corresponding experimental condition. No specific version of the reference genome is required, as users provide both the reference sequence and its annotations. These can be obtained from public resources such as UCSC^36,37^ and RepeatMasker, or generated de novo using workflows that combine RepeatModeler^38,39^ and RepeatMasker. Generating new custom annotations could improve repeat identification, and they are compatible with the PERREO pipeline if provided in GTF format.

### Software and computational environment

The PERREO pipeline (v1.0.0) was executed under Conda (v24.7.1) on a Linux-based cluster (Rocky Linux 8.5), using R (v4.2.3) and Python (v3.11.0). A complete specification of all software dependencies and their versions is provided in the Conda environment files deposited together with the PERREO code (see Code availability). Additionally, TEtranscripts v2.2.3 and Salmon v1.10.3 were used.

### PERREO for long-read data

To analyze direct RNA data generated with Oxford Nanopore technology, a suitable repeat-aware reference must be prepared prior to running the pipeline. In our case, we downloaded the RepeatMasker track for the CHM13-T2T v2.0 assembly from the UCSC Genome Browser in FASTA format, together with the corresponding RepeatMasker annotations made available by the T2T consortium.

PERREO expects basecalled reads in FASTQ format, assuming that basecalling has been performed with dedicated software such as Dorado^40^. Before launching the pipeline, read quality is evaluated with NanoPlot^41^, which helps users select the most appropriate parameter settings. Reads are then aligned to the reference genome using minimap2, with settings optimized for spliced direct RNA data (-ax splice -uf -k14 -p 0.8-N 100).

### PERREO for short-read data

For short-read datasets, raw FASTQ files are first trimmed with cutadapt^42^ using the parameters -q 30,30 -m 16. If adapter sequences are not explicitly specified, trimming is performed without adapter removal. When the sequencing technology is known to introduce artificial GC nucleotides at the 3′ end of reads, users can invoke an additional trimming step that calls the trimGC.py^43^ script to remove these artefactual bases.

After trimming, read quality is assessed using FastQC. Users can review FastQC reports once the run is complete to confirm that the data meet their quality expectations. Read alignment is performed with parameters specifically tuned to retain multimapping reads, as repRNAs can map to multiple genomic loci. In our tests, setting both --outFilterMultimapNmax and --winAnchorMultimapNmax to 500 provided a suitable balance between sensitivity and specificity for repeat mapping, while allowing up to 5% mismatches by default to reduce spurious alignments.

### Downstream analysis

#### Quantification

Downstream steps are shared between short-read- and long-read-based analyses. Quantification is performed with the featureCounts function in R, enabling the countMultiMappingReads and fraction options to appropriately handle multimappers, and specifying the library strandedness as defined in the sample sheet.

#### Differential expression analysis (DEA)

Differential expression analysis is carried out using either edgeR or DESeq2, without restriction on the number of experimental conditions, as the design can accommodate multiple groups. For short-read data, the count matrix is filtered to retain only features with non-zero counts across samples. For long-read data, we allow up to 10% zero counts per feature, reflecting the higher sparsity typical of long-read-derived count tables.

To reduce unwanted technical variation, users may optionally apply the RUVg function from the RUVSeq package^44^. This method models unobserved factors contributing to unwanted variability, controlled by the parameter k (default k = 2), which can be adjusted by providing the -k argument.

PERREO also allows the user to specify a batch-effect factor, which must be included as an additional column in the sample sheet (for example, the hospital or sequencing center of origin).

Data normalization depends on the chosen DEA method: DESeq2-based analyses use variance-stabilizing transformation (vst), whereas edgeR-based analyses use the TMM algorithm. By default, repRNAs are considered differentially expressed when ∣log_2_FC∣>1 and adjusted p-value < 0.05. These thresholds can be customized using the - log2FC and -FDR arguments. The workflow additionally generates a set of summary tables and graphical outputs to facilitate interpretation.

#### Transcriptome assembly

To explore transcript diversity and identify novel transcripts not present in the original annotation, PERREO integrates StringTie2^45^. This step produces an assembled transcriptome file for each sample, enabling the detection of new repeat-containing isoforms and hybrid transcripts.

#### Coexpression analysis

To investigate coordinated expression patterns, the pipeline implements coexpression network analysis using the WGCNA R package^46^. The workflow returns a complete list of repeat features and their module assignments, as well as detailed information for the three modules showing the strongest correlation with any experimental condition. These outputs can be directly imported into Cytoscape^47^ to build and visualize coexpression networks.

#### Prediction model performance

Machine-learning algorithms are used to evaluate the predictive potential of repRNAs as biomarkers. PERREO includes two complementary approaches: Random Forest, a non-linear tree-based ensemble, and GLMnet, a regularized linear model. Both models are trained using the caret R package^48^, allowing users to compare how linear versus non-linear decision functions perform on their specific dataset (Supplementary Note 2.3).

### Portability

PERREO offers different execution modes to accommodate distinct levels of computational expertise. A user-friendly graphical interface allows non-programmers to configure and run the pipeline or to automatically generate the corresponding command-line code for execution in a terminal. The script to execute the app is also included in our GitHub repository.

All required tools and R/Python packages can be installed via a dedicated Conda environment defined in a YAML file provided in the project’s GitHub repository. By default, the pipeline runs using 8 threads and 32 GB of RAM and is designed to be fault-tolerant: if an error occurs, users can rerun the command after addressing the issue, and PERREO will detect existing output files, skip completed steps, and resume the workflow from the appropriate point.

## Data availability

The datasets analyzed in this study can be found in the Singapore Nanopore Expression Project AWS Open Data link (https://registry.opendata.aws/sgnex/) and Gene Expression Omnibus (https://www.ncbi.nlm.nih.gov/geo/) under the following accession codes: GSE147352, GSE174302, and GSE228512, GSE242689, and GSE117387. Pre-built annotations file of repetitive elements for T2T-CHM13 reference genome was downloaded from the dropbox link included in MGH lab website (https://www.mghlab.org/software/tetranscripts).

## Code availability

All the required scripts and instructions are stored in our GitHub repository: https://github.com/DGCLab/PERREO-Pipeline.

## Acknowledgments

Thanks to Diana Aguilar Morantés lab and our group members for helpful discussions.

## Funding

AECC-Estratégico2023 (PRYES235086GOME)

PID2022-137280OB-I00 CNS2023-143939

Andalusian Regional Government, CSyC 2024 - Proyectos (RPS 24664), PI-0161-2024

## Author contributions

Conceptualisation: DGC, FRM

Methodology: DGC, FRM, MMS

Investigation: DGC, FRM,

Supervision: DGC

Writing—original draft: FRM, DGC

Writing—review & editing: FRM, DGC

## Data and materials availability

All data, code, and materials used in the analyses must be available in some form to any researcher for purposes of reproducing or extending the analyses.

## Extended Data legends

**Extended Data Fig. 1:**
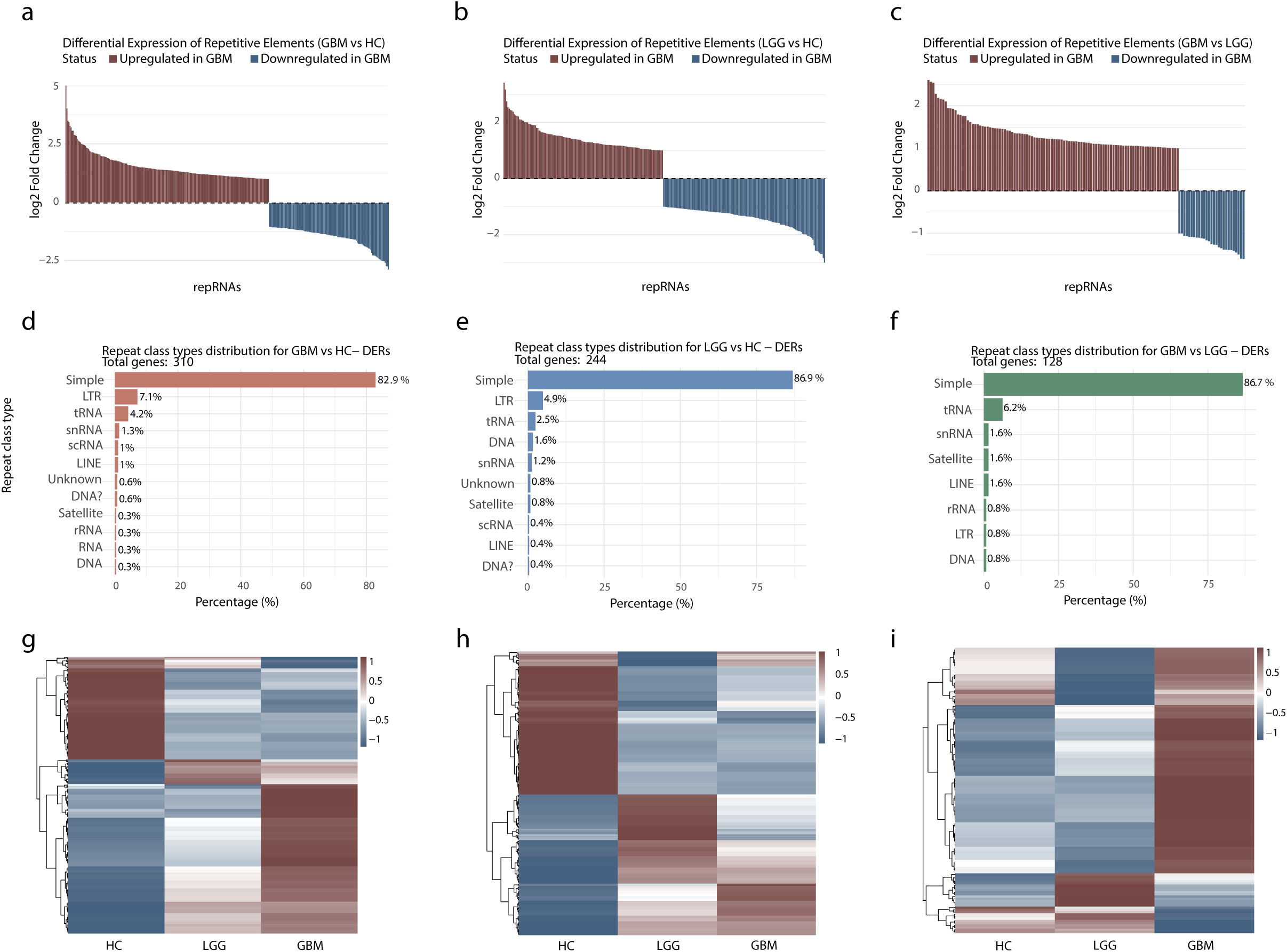
Repeat RNAs analysis in glioblastoma and low-grade glioma using GRCh38 assembly. **a, b, c,** Barplot of differentially expressed features in GBM versus HC **(a)**, LGG versus HC **(b)** and GBM versus LGG **(c)** contrasts. **d, e, f,** Most represented repetitive classes in GBM versus HC **(d)**, LGG versus HC **(e)** and GBM versus LGG **(f)** DERs; expression heatmap of DERs in GBM versus HC **(g)**, LGG versus HC **(h)** and GBM versus LGG **(i)** contrasts.

**Extended Data Fig. 2:**
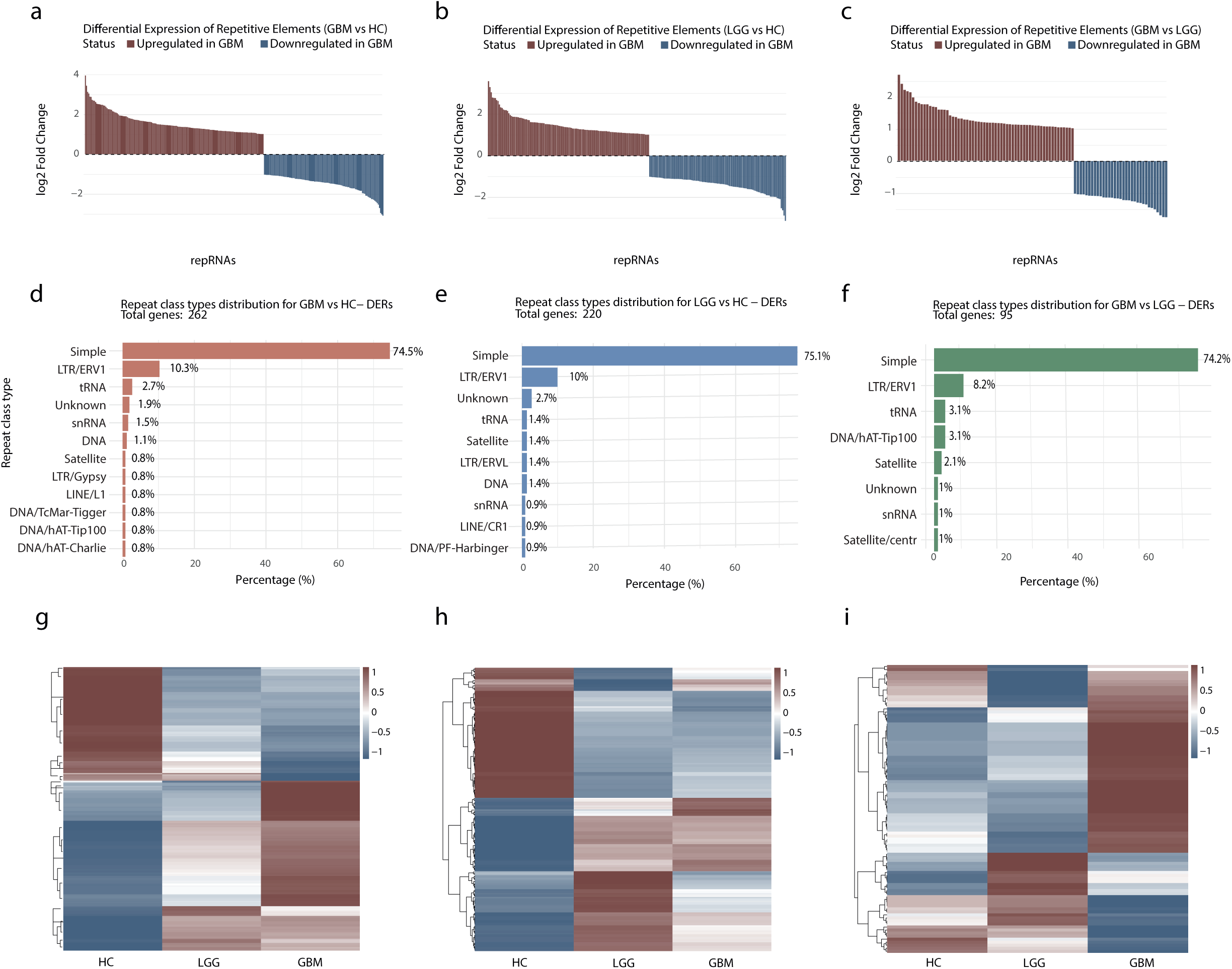
Repeat RNAs analysis in glioblastoma and low-grade glioma using T2T assembly. **a, b, c,** Barplot of differentially expressed features in GBM versus HC **(a)**, LGG versus HC **(b)** and GBM versus LGG **(c)** contrasts. **d, e, f,** Most represented repetitive classes in GBM versus HC **(d)**, LGG versus HC **(e)** and GBM versus LGG **(f)** DERs; expression heatmap of DERs in GBM versus HC **(g)**, LGG versus HC **(h)** and GBM versus LGG **(i)** contrasts.

**Extended Data Fig. 3:**
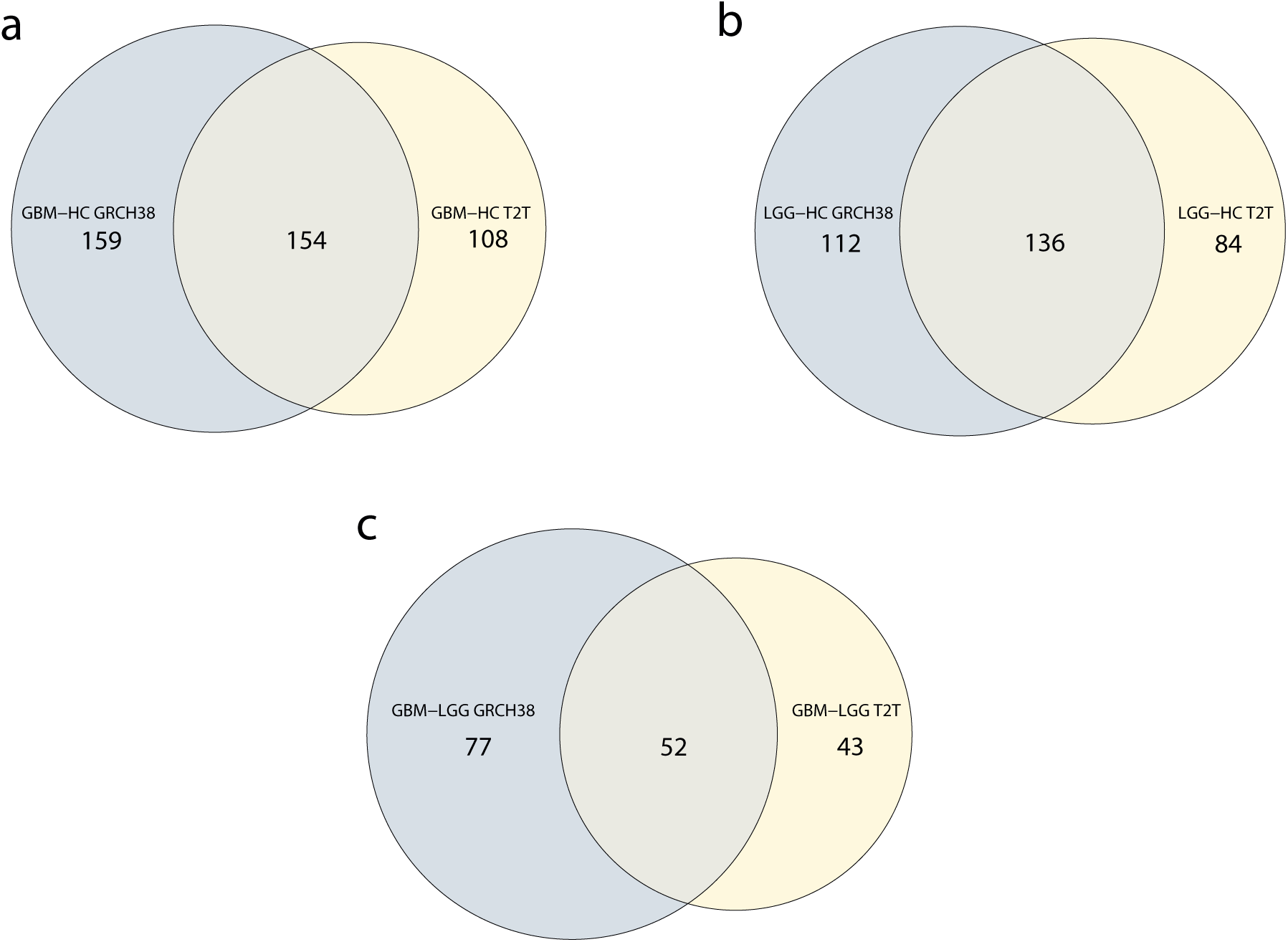
Common and unique differentially expressed features in glioblastoma and low-grade glioma using GRCh38 and T2T references. **a, b, c,** Venn diagrams comparing DERs obtained using GRCh38 and T2T reference genomes for each contrast: GBM versus HC **(a)**, LGG versus HC **(b)**, and GBM versus LGG **(c)**.

**Extended Data Fig. 4:**
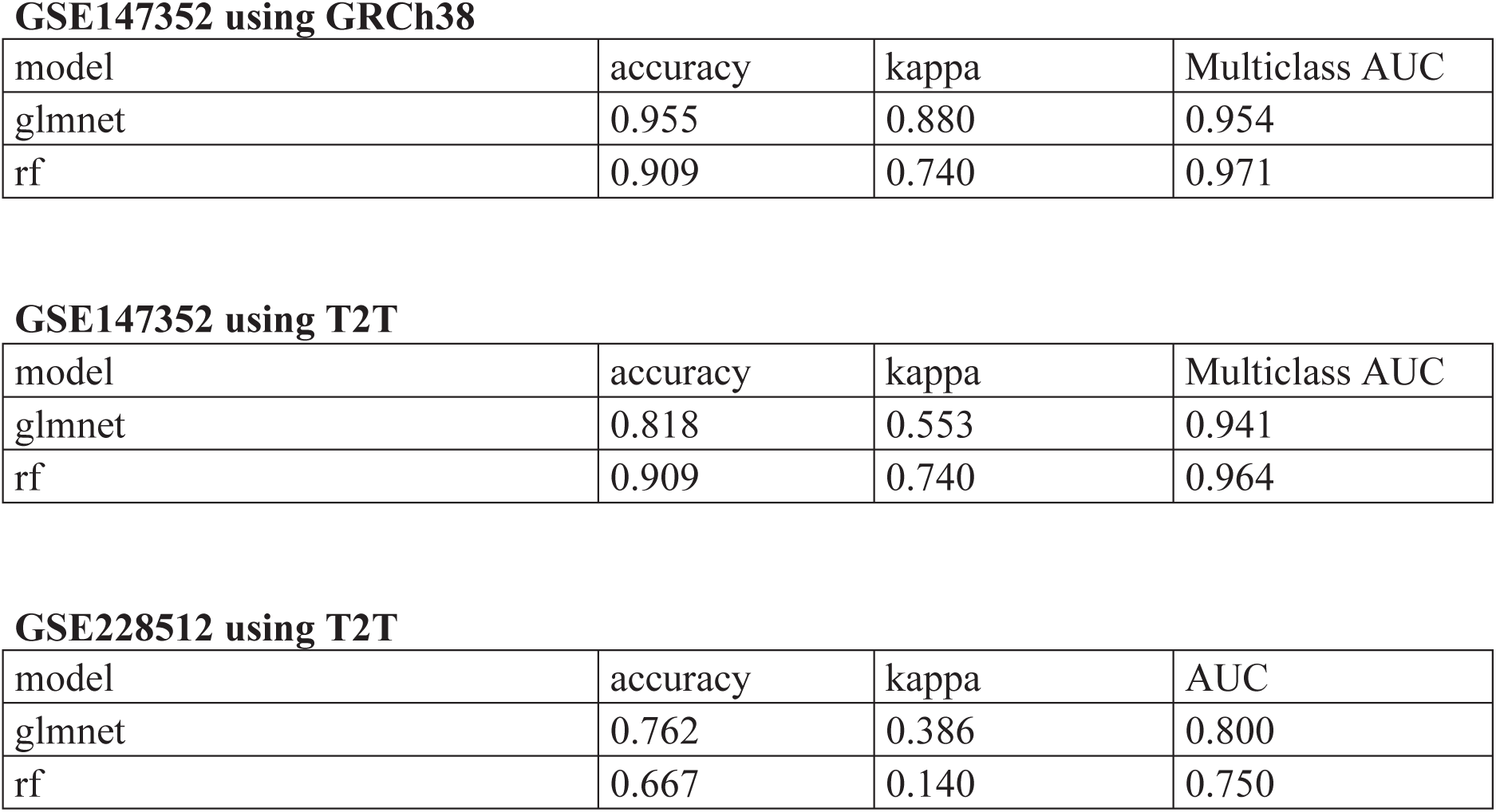
GLMnet and Random Forest prediction algorithms performance on GSE147352 and GSE228512 datasets. Prediction models results obtained from the analysis of GSE147352 and GSE228512 datasets.

**Extended Data Fig. 5:**
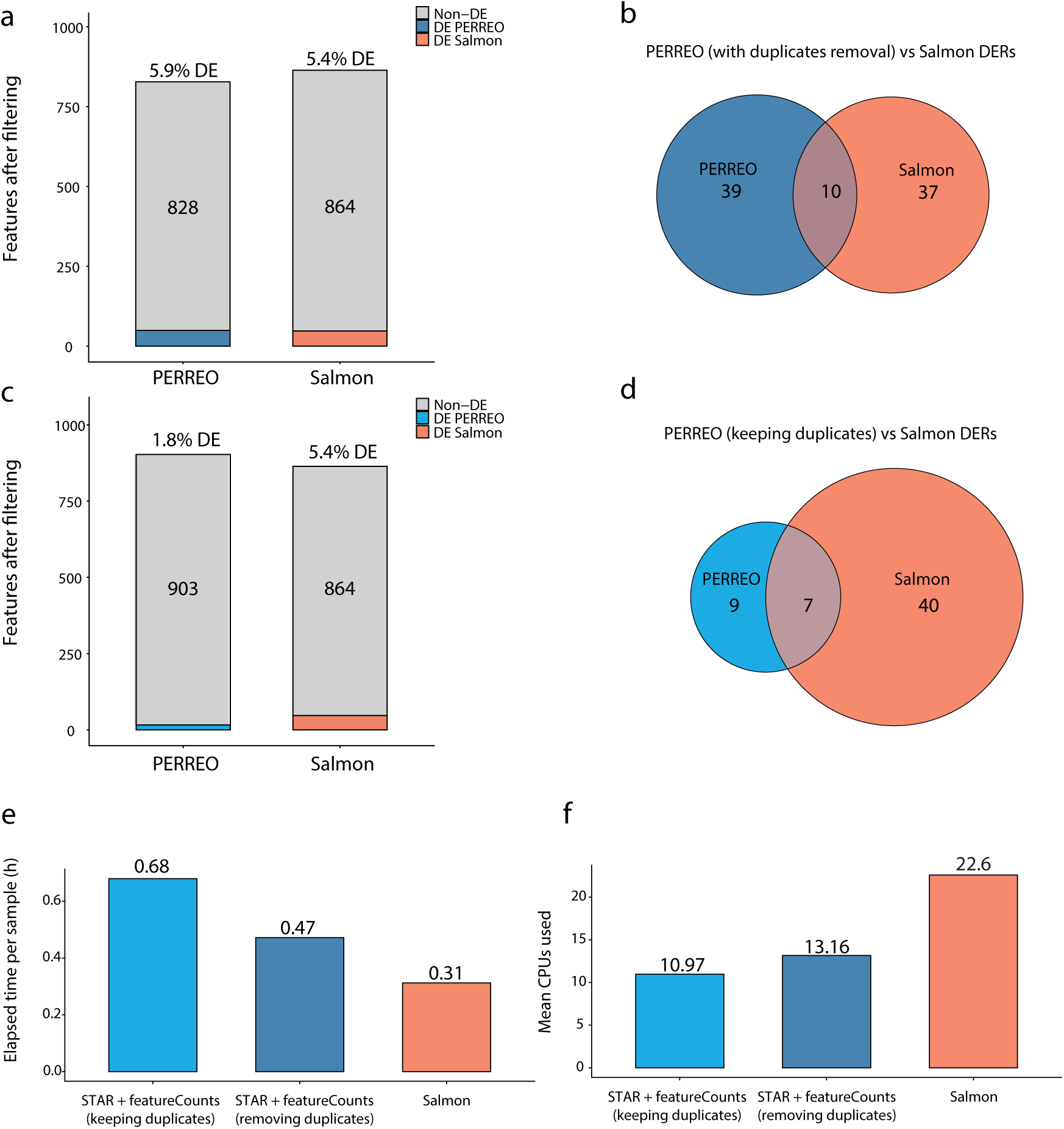
Comparing PERREO results against results derived from running Salmon pseudoaligner on GSE174302 dataset. **a, b,** Comparison of results obtained with Salmon and PERREO (the latter removing duplicate reads). **a,** Barplot showing total repeat identifications and corresponding DERs using PERREO discarding duplicates and Salmon pseudoaligner. **b,** Venn diagram representing common and specific DERs using previously mentioned approaches. **c,d,** Comparison of results obtained with Salmon and PERREO (the latter retaining duplicate reads). **c,** Barplot showing total repeat identifications and corresponding DERs using PERREO keeping duplicates and Salmon pseudoaligner. **d,** Venn diagram representing common and specific DERs recently mentioned approaches. **e, f,** Barplots containing the information about the performance of Salmon and PERREO (keeping and removing duplicates) in terms of elapsed time **(e)** and mean CPUs used **(f)**.

**Extended Data Fig. 6:**
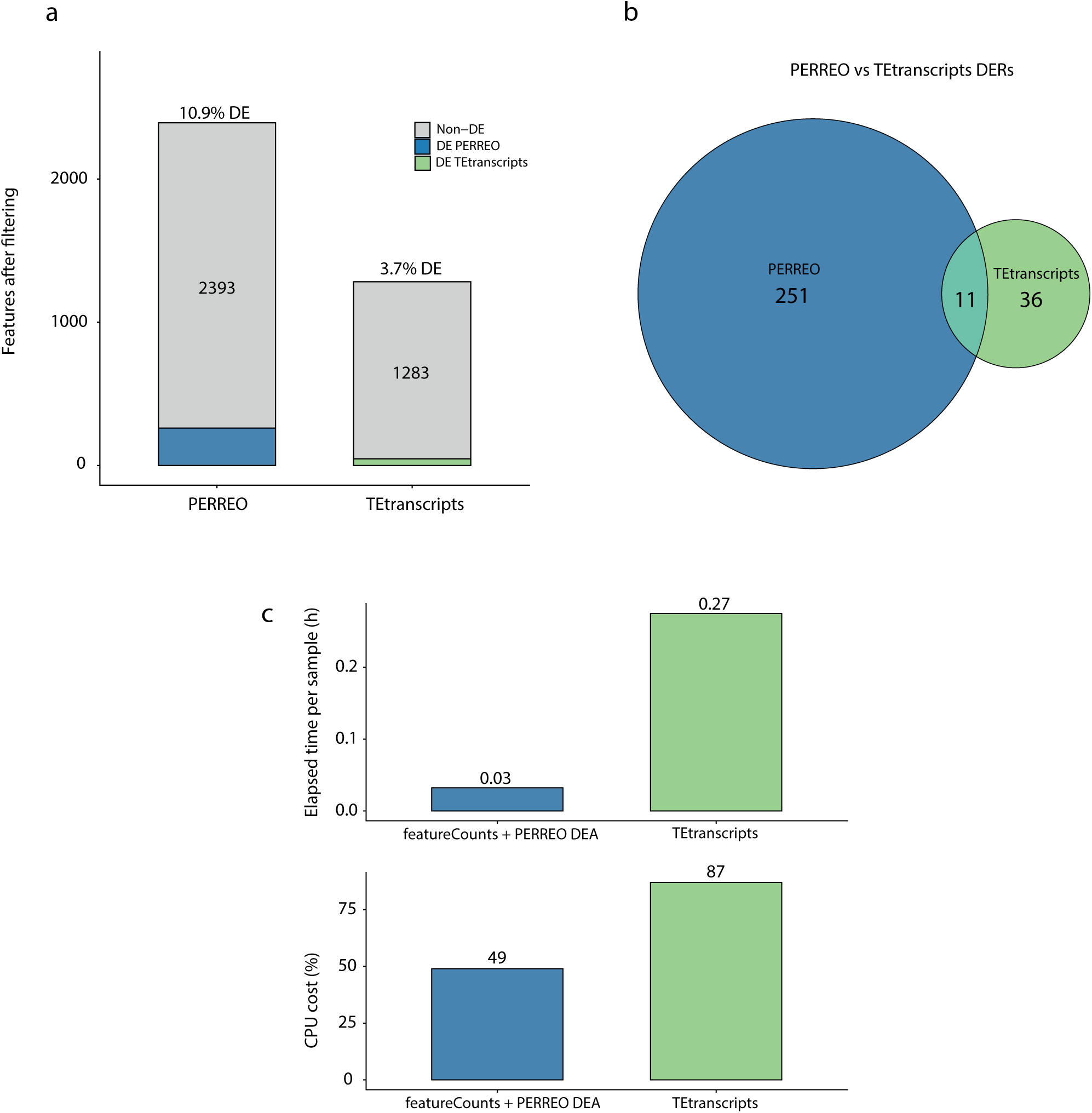
Benchmarking PERREO against TEtranscripts TE-based software using GSE147352 dataset as subject. **a,** Bar plot showing the number of total identified features after data filtering using PERREO and TEtranscripts, as well as the number of DERs identified by both pipelines. **b,** Venn diagram showing common and method-specific DERs for PERREO and TEtranscripts. **c,** Summary of the performance of featureCounts + PERREO DEA and TEtranscripts in terms of elapsed time per sample and CPU usage (%).

## REFERENCES

1. Espinosa, E., Bautista, R., Larrosa, R. & Plata, O. Advancements in long-read genome sequencing technologies and algorithms. Genomics 116, 110842 (2024).

2. Criscione, S. W., Zhang, Y., Thompson, W., Sedivy, J. M. & Neretti, N. Transcriptional landscape of repetitive elements in normal and cancer human cells. BMC Genomics 15, 583 (2014).

3. Jin, Y., Tam, O. H., Paniagua, E. & Hammell, M. TEtranscripts: a package for including transposable elements in differential expression analysis of RNA-seq datasets. Bioinformatics 31, 3593–3599 (2015).

4. Yang, W. R., Ardeljan, D., Pacyna, C. N., Payer, L. M. & Burns, K. H. SQuIRE reveals locus-specific regulation of interspersed repeat expression. Nucleic Acids Res. 47, e27–e27 (2019).

5. Hoyt, S. J. et al. From telomere to telomere: The transcriptional and epigenetic state of human repeat elements. Science 376, eabk3112 (2022).

6. Reggiardo, R. E. et al. Profiling of repetitive RNA sequences in the blood plasma of patients with cancer. *Nat*. Biomed. Eng. 7, 1627–1635 (2023).

7. Cherkasova, E. et al. Detection of an Immunogenic HERV-E Envelope with Selective Expression in Clear Cell Kidney Cancer. Cancer Res. 76, 2177–2185 (2016).

8. Harris, C. R. et al. Association of Nuclear Localization of a Long Interspersed Nuclear Element-1 Protein in Breast Tumors with Poor Prognostic Outcomes. Genes Cancer 1, 115–124 (2010).

9. Meng, X. et al. TE-SCALE: a comprehensive database for exploring transposable element expression across human cancers at single-cell resolution. Nucleic Acids Res. 54, D1658–D1671 (2026).

10. De Luca, C. et al. Enhanced expression of LINE-1-encoded ORF2 protein in early stages of colon and prostate transformation. Oncotarget 7, 4048–4061 (2016).

11. Zadran, B. et al. Impact of retrotransposon protein L1 ORF1p expression on oncogenic pathways in hepatocellular carcinoma: the role of cytoplasmic PIN1 upregulation. Br. J. Cancer 128, 1236–1248 (2023).

12. Taylor, M. S. et al. Ultrasensitive Detection of Circulating LINE-1 ORF1p as a Specific Multicancer Biomarker. Cancer Discov. 13, 2532–2547 (2023).

13. Dobin, A. et al. STAR: ultrafast universal RNA-seq aligner. Bioinformatics 29, 15–21 (2013).

14. Liao, Y., Smyth, G. K. & Shi, W. featureCounts: an efficient general purpose program for assigning sequence reads to genomic features. Bioinformatics 30, 923–930 (2014).

15. Li, H. Minimap2: pairwise alignment for nucleotide sequences. Bioinformatics 34, 3094–3100 (2018).

16. Robinson, M. D., McCarthy, D. J. & Smyth, G. K. edgeR : a Bioconductor package for differential expression analysis of digital gene expression data. Bioinformatics 26, 139–140 (2010).

17. Love, M. I., Huber, W. & Anders, S. Moderated estimation of fold change and dispersion for RNA-seq data with DESeq2. Genome Biol. 15, 550 (2014).

18. Ewels, P., Magnusson, M., Lundin, S. & Käller, M. MultiQC: summarize analysis results for multiple tools and samples in a single report. Bioinformatics 32, 3047–3048 (2016).

19. Andrew, S. FastQC: A Quality Control Tool for High Throughput Sequence Data. Available online at: http://www.bioinformatics.babraham.ac.uk/projects/fastqc/. (2010).

20. Bao, Y., Zhang, D., Guo, H. & Ma, W. Beyond blood: Advancing the frontiers of liquid biopsy in oncology and personalized medicine. Cancer Sci. 115, 1060–1072 (2024).

21. Chen, Y. et al. A systematic benchmark of Nanopore long-read RNA sequencing for transcript-level analysis in human cell lines. Nat. Methods 22, 801–812 (2025).

22. Xiao-Jie, L., Hui-Ying, X., Qi, X., Jiang, X. & Shi-Jie, M. LINE-1 in cancer: multifaceted functions and potential clinical implications. Genet. Med. 18, 431–439 (2016).

23. Wissing, S. et al. Reprogramming somatic cells into iPS cells activates LINE-1 retroelement mobility. Hum. Mol. Genet. 21, 208–218 (2012).

24. Liu, Q. & Li, J. Computational tools for tandem repeat detection using long-read sequencing. Brief. Bioinform. 27, bbag031 (2026).

25. Haile, S. et al. Adaptable and comprehensive approaches for long-read nanopore sequencing of polyadenylated and non-polyadenylated RNAs. Front. Genet. 15, 1466338 (2024).

26. Begik, O. et al. Nano3P-seq: transcriptome-wide analysis of gene expression and tail dynamics using end-capture nanopore cDNA sequencing. Nat. Methods 20, 75–85 (2023).

27. Kim, D., Paggi, J. M., Park, C., Bennett, C. & Salzberg, S. L. Graph-based genome alignment and genotyping with HISAT2 and HISAT-genotype. Nat. Biotechnol. 37, 907–915 (2019).

28. Patro, R., Duggal, G., Love, M. I., Irizarry, R. A. & Kingsford, C. Salmon provides fast and bias-aware quantification of transcript expression. Nat. Methods 14, 417–419 (2017).

29. Smit, A. F. A., Hubley, R. & Green, P. RepeatMasker Open-4.0. (2013).

30. Do, C. B. & Batzoglou, S. What is the expectation maximization algorithm? Nat. Biotechnol. 26, 897–899 (2008).

31. Deschamps-Francoeur, G., Simoneau, J. & Scott, M. S. Handling multi-mapped reads in RNA-seq. Comput. Struct. Biotechnol. J. 18, 1569–1576 (2020).

32. Kan, L. K. et al. Potential biomarkers and challenges in glioma diagnosis, therapy and prognosis. *BMJ Neurol*. Open 2, e000069 (2020).

33. Nogalski, M. T. & Shenk, T. HSATII RNA is induced via a noncanonical ATM-regulated DNA damage response pathway and promotes tumor cell proliferation and movement. Proc. Natl. Acad. Sci. 117, 31891–31901 (2020).

34. Shah, A. H. et al. Differential expression of an endogenous retroviral element [HERV-K(HML-6)] is associated with reduced survival in glioblastoma patients. Sci. Rep. 12, 6902 (2022).

35. You, E. et al. Disruption of cellular plasticity by repeat RNAs in human pancreatic cancer. Cell 187, 7232–7247.e23 (2024).

36. Hinrichs, A. S. et al. The UCSC Genome Browser Database: update 2006. Nucleic Acids Res. 34, (2006).

37. Karolchik, D. et al. The UCSC Table Browser data retrieval tool. Nucleic Acids Res. 32, (2004).

38. Smit, A. F. A. & Hubley, R. RepeatModeler Open-1.0. (2008).

39. Flynn, J. M. et al. RepeatModeler2 for automated genomic discovery of transposable element families. Proc. Natl. Acad. Sci. 117, 9451–9457 (2020).

40. Oxford Nanopore Technologies. Dorado basecaller.

41. De Coster, W., D’Hert, S., Schultz, D. T., Cruts, M. & Van Broeckhoven, C. NanoPack: visualizing and processing long-read sequencing data. Bioinformatics 34, 2666–2669 (2018).

42. Martin, M. Cutadapt removes adapter sequences from high-throughput sequencing reads. 17, 10–12 (2011).

43. Chen, S. et al. Cancer type classification using plasma cell-free RNAs derived from human and microbes. eLife 11, e75181 (2022).

44. Risso, D., Ngai, J., Speed, T. & Dudoit, S. Normalization of RNA-seq data using factor analysis of control genes or samples. 32, 896–902 (2014).

45. Kovaka, S. et al. Transcriptome assembly from long-read RNA-seq alignments with StringTie2. Genome Biol. 20, 278 (2019).

46. Langfelder, P. & Horvath, S. WGCNA: an R package for weighted correlation network analysis. BMC Bioinformatics 9, 559 (2008).

47. Shannon, P. et al. Cytoscape: A Software Environment for Integrated Models of Biomolecular Interaction Networks. Genome Res. 13, 2498–2504 (2003).

48. Kuhn, M. Building Predictive Models in *R* Using the **caret** Package. J. Stat. Softw. 28, (2008).

