## Supplementary Notes for "PERREO: An integrated pipeline for repetitive elements analysis enables the repeatome expression profiling in cancer"

### Supplementary information

#### Index

#### Supplementary Note 1: Running PERREO on user-friendly interface

Bioinformatics analyses can be run either directly from a terminal or through the graphical interface that we developed. Within this interface, the “Usage” section provides detailed instructions for running the pipeline in all supported modes (SR-SE, SR-PE, and LR). For each mode, users can specify parameter values and select the options that best fit their experimental design. There are two main options for execution: running the pipeline directly from the interface or copying the automatically generated command and pasting it into a terminal.

Quality control reports generated by FastQC and MultiQC can be visualized within the interface, allowing users to adjust parameters based on these quality metrics. The interface continuously updates a command line that can be used to run the pipeline in the user’s terminal. To ensure correct execution, all files associated with the interface must be stored in the same top-level working directory as the pipeline, so that all required scripts and configuration files are accessible.

#### Supplementary Note 2: Extended workflow and implementation details

##### *2.1. Coexpression analysis with WGCNA*

We implemented a workflow to identify coexpression modules and relationships among repeat RNAs that display similar behaviour across samples. The program automatically selects the lowest soft-thresholding power that yields a scale-free topology with $R^2 > 0.9$. Once the network has been constructed from the expression matrix, several visualizations are generated, including a module dendrogram and a module–condition correlation heatmap. In addition, the workflow exports two text files containing node and edge information for the network, respectively, as well as separate text files listing the features assigned to each identified module.

##### *2.2. Transcriptome assembly to further characterize the emergence of novel transcripts from repetitive regions*

Transcriptome assembly with StringTie enables the identification of chimeric RNAs and transcripts synthesized from non-coding and repetitive regions, providing a complementary view to feature-level repeat quantification. Genomic instability and widespread transcriptional deregulation in cancer can give rise to gene fusions and repeat-derived transcripts that are rarely observed under normal conditions, and PERREO’s assembly module helps pinpoint such novel repeat-containing isoforms and their genomic context. Beyond cataloguing these events, this analysis offers a starting point to investigate the potential regulatory roles of novel repeat-derived transcripts and their contribution to shaping the tumor microenvironment. Molecular alterations at nucleotide level, like insertions and deletions, can also be analyzed in aligned reads as well as spliced junctions and alternative splicing products. Additionally, StringTie results can be used to perform subsequent downstream analysis with Salmon.

##### *2.3. Prediction models' design*

The caret R package was used to build prediction models based on Random Forest and GLMnet algorithms. Parallelization was enabled according to the number of threads specified in the code. Both the number of folds and the number of complete sets of folds for k-fold cross-validation were set to 5, using the "repeatedcv" resampling method. Depending on the number of experimental conditions in the design, model performance was summarized using either a two-class summary or a multiclass summary function.

The source code was designed to balance and optimize the distribution of samples across conditions. When the ratio between the number of samples in the majority and minority classes was greater than 0.7, no resampling was applied ("none"). Conversely, when the minority class contained fewer than 20 samples, downsampling ("down") was used. The createDataPartition function was then applied to split the data into training and test sets, with 80% of the samples automatically assigned to the training set.

The pipeline generates summary tables with performance metrics for each prediction model, as well as plots showing the AUC values for the Random Forest and GLMnet models.

#### Supplementary Note 3: Extended repeatome analyses across datasets

##### *3.1. Repeatome profile of dogs after immunotherapy treatment*

We tested PERREO SR-SE mode with the dataset stored in GSE242689. Bulk RNA-seq experiments were conducted on samples obtained from dogs with canine mammary cancer that were treated with a specific type of immunotherapy based on cowpea mosaic virus. We compared the expression profiles of these samples before and after treatment to determine the specific effect of this therapy on their repeatome expression profile.

We used Dog10K_Boxer_Tasha assembly from Ensembl as reference with its corresponding annotations. The repeats annotations were downloaded from RepeatMasker track in UCSC table browser. We ran the pipeline indicating the parameters -trimming simple -polya polya -batch no -method DESeq2 -remove_duplicates yes. Only two features were differentially expressed after statistical analysis, that were LTR53-int and Eutr10. Both were downregulated in post-treatment conditions with respect to pre-treatment conditions.

##### *3.2. Repeats RNAs expression in mice mammary tumors*

The GSE117387 dataset contains mammary tissue samples derived from mammary tumors and normal tissue. Single-end sequencing was performed, so we ran PERREO in SR-SE mode using the GRCm39/mm39 genome from UCSC as reference, together with the corresponding gene annotations and repeat annotations from the RepeatMasker track. The pipeline was executed with the options -trimming simple -remove_duplicates false -batch no -method edgeR.

In this dataset, the roles of Wwox and Trp53 were investigated using genetically engineered mouse models. Among the 10 tumor samples, 2 carried Trp53 mutations, 4 carried Wwox mutations, and 4 carried mutations in both genes. Among the 6 normal mammary epithelial cell samples, 3 were obtained from wild-type mice and 3 from Wwox-knockout mice.

Preliminary analyses showed that PCA clearly separated tumor samples from normal epithelial cell samples but did not reveal marked differences between genotypes within the same experimental condition. Therefore, we focused on analysing repeatome expression patterns in tumors versus normal tissue, independently of genotype, and observed that many features displayed opposite expression profiles between the two groups.

Overall, we detected a clear increase in mean repetitive element expression in tumors, and PCA robustly separated the two experimental conditions. Using the default differential expression thresholds in PERREO, 942 features were classified as differentially expressed.

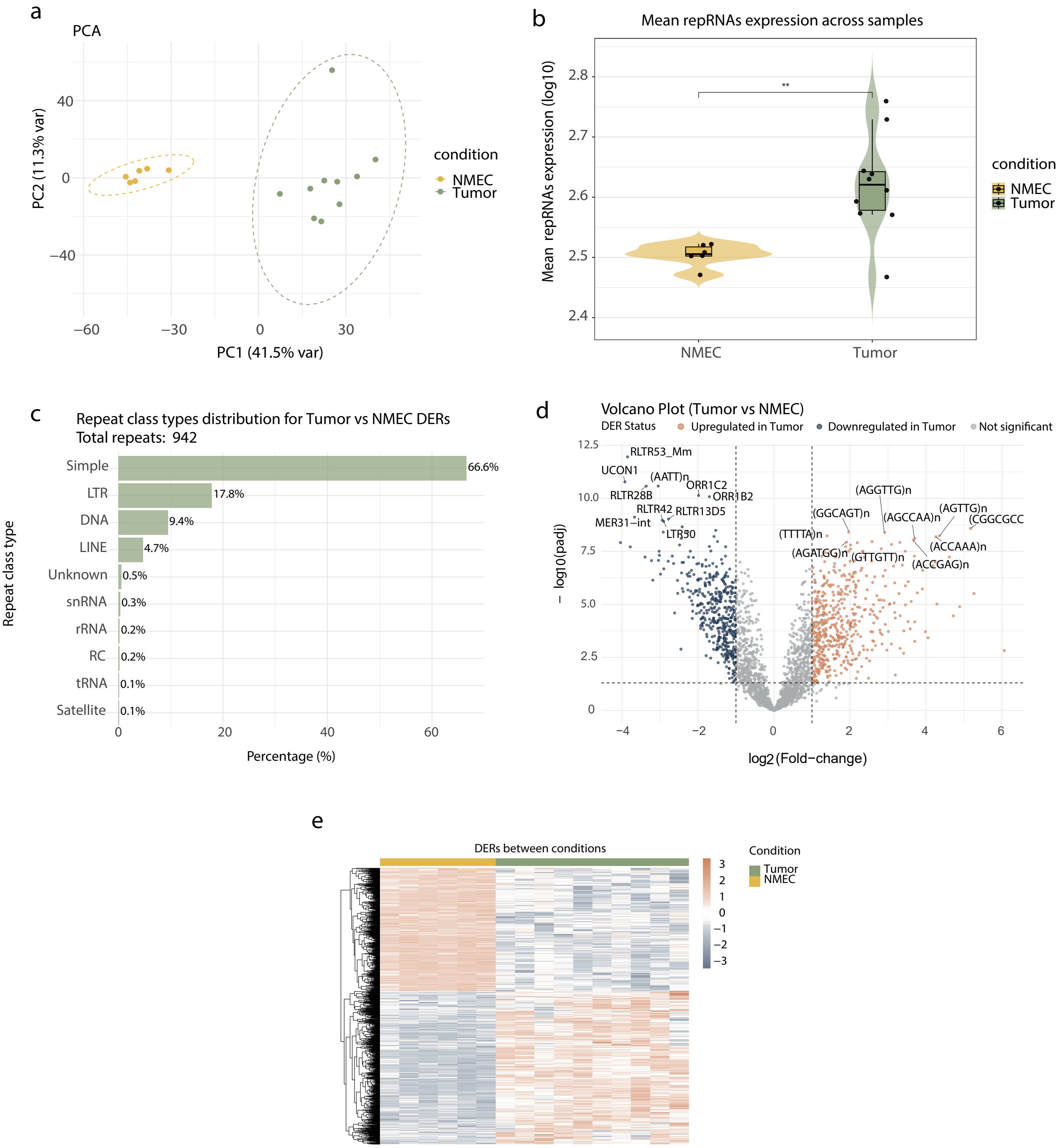

**Figure 1. a,** PCA displaying NMEC and Tumor data distribution. **b,** Violin plot showing the median, upper and lower quartiles of repeat RNAs expression in NMEC and tumors. Significance level was calculated with Wilcoxon test, ****P < 0.0001, ***P < 0.001, **P < 0.01, *P < 0.05, 'ns' P > 0.05. **c,** Repeat classes distribution of DERs in Tumor versus NMEC contrast. **d,** Volcano plot showing upregulated and downregulated repRNAs in Tumor versus NMEC contrast in orange and blue, respectively. **e,** Expression heatmap of DERs.

#### Supplementary Note 4: Benchmarking alternative alignment and quantification workflows

##### *4.1. Evaluation of alignment–quantification strategies for repeat RNA analysis*

As described in the manuscript, we tried to compare three different workflows: HISAT2, STAR and Salmon. We analyzed transcriptomics data obtained from oesophagus cancer patients' plasma (GSE174302) and we compared their performance with PERREO workflow constituted by cutadapt, STAR aligner and featureCounts. Pipelines were run remotely in a computing cluster using 32 threads and 130 RAM GB for the process in computing nodes with 34 threads and 150 RAM GB capacity.

We ran HISAT2 with the parameters -k 10 --seed 42 and subsequent on-the-fly sorting via samtools sort, several samples reproducibly failed at the alignment step, with errors such as truncated BAM files and SAM records with mismatched SEQ and QUAL lengths at identical computational resources. These issues, which likely reflect a combination of the high multimapping burden of repeat-derived reads and the specific parameterization used, prevented us from obtaining a complete and reliable HISAT2-based count matrix. In contrast, the STAR + MarkDuplicates + featureCounts workflow ran successfully for all samples under the same conditions and therefore was used as the primary mapping–counting strategy in this study. On the other hand, we would have increased the –k parameter to report as many multimapped RNA copies as STAR, but we were not able to do so due to technical issues.

Additionally, Salmon was also run, configuring the index with --keepDuplicates and running salmon quant with --validateMappings, --numBootstraps 30, --seqBias, --gcBias, and –posBias and using as reference a repeats transcripts fasta with unique sequences file obtained from UCSC browser. In line with the developers' recommendations against removing duplicates in the absence of UMIs, we kept all reads and used the same statistical thresholds as in the STAR + featureCounts workflow. Under these conditions, Salmon detected 47 differentially expressed features, 10 of which overlapped those identified by STAR + featureCounts. Overall, Salmon provided faster read processing than genome-based alignment workflows, but at the cost of not retaining explicit genomic coordinates, which limits direct locus-level interpretation of repeat expression.

To obtain results that were maximally comparable to those derived from Salmon, we ran STAR followed by featureCounts without removing duplicate reads on the same set of 33 samples. This workflow took 22 hours and 24 minutes and the mean number of CPUs used during the task were 10.97 (Table 1). Furthermore, we ran DEA again with the resulting count matrix, and we found only 16 DERs, 7 of which were also identified as DERs in the Salmon-based workflow.

| | STAR + featureCounts (removing duplicates) | STAR + featureCounts (keeping duplicates) | Salmon |
|---|---|---|---|
| Elapsed time (h:mm:ss) | 16:02:30 | 22:24:43 | 10:37:08 |
| Threads | 32 | 32 | 32 |
| CPU cost (%) | 1316 | 1097 | 2260 |
| Mean CPUs used | 13.16 | 10.97 | 22.6 |
| N (samples) | 34 | 33 | 34 |

**Table 1.** Computational resource usage results for STAR + featureCounts and Salmon pipelines.

##### *4.2. Comparison with TEtranscripts, a TE-focused workflow*

We conducted DEA with TEtranscripts software, using identical thresholds (FDR < 0.01, |log2FC| > 1) and pre-built curated annotations for the T2T genome, from MGH lab website (https://www.mghlab.org/software/tetranscripts).

We ran the program with this configuration to reply PERREO analysis as possible: --stranded reverse --mode multi --padj 0.01 --foldchange 1. TEtranscripts -mode multi option is designed to deal with multimapped reads using the same approach as PERREO, dividing their counts into the number of sites where these reads are mapped. However, it additionally uses EM algorithm to estimate the probability of each read to be transcribed from each region it mapped to refine the fraction of each read.

TEtranscripts analyze transcriptomics data from alignment BAM files, separating traditional genomic annotations features and repetitive elements features. In this way, we compared the performance of featureCounts and DEA scripts designed for PERREO pipeline with respect to the performance of TEtranscripts complete workflow using the BAM files obtained from the previously mentioned GSE147352 dataset.

TEtranscripts did not provide any option to indicate the number of threads to be used in the process, while PERREO was run using 14 threads in our local computer. Additionally, we could not include LGG samples in TEtranscripts pipeline as it is designed to compare samples that belong to two specific experimental conditions.

TEtranscripts and PERREO results are not directly comparable, as the higher algorithmic complexity of TEtranscripts entails increased CPU usage and runtime. TEtranscripts is specifically designed to handle multimapping reads and reduce bias in young TE subfamilies by grouping similar sequences into subfamilies, which is highly beneficial for characterizing global patterns across repetitive element families. In contrast, PERREO distributes multimapping reads fractionally because it is not focused on pinpointing the specific genomic copies that are expressed, but rather on quantifying the expression level of each repeat, which is more suitable for marker-oriented analyses.

Differences between both approaches were expected since PERREO applies stringent count-based filtering before differential analysis. Even so, PERREO retained 2,393 repeat features, compared to only 1,283 in the TEtranscripts-derived matrix, a discrepancy likely arising from differences in annotation files. TEtranscripts detected only 47 features as differentially expressed in GBM-HC comparisons, 11 of which overlapped with PERREO results. Among these, four were downregulated and seven upregulated, showing consistent expression trends across both analyses.

In terms of computational performance, TEtranscripts required nearly 27 hours to complete, using 87% of CPU resources. By contrast, PERREO's featureCounts and differential expression steps finished in only 3 hours and 42 minutes, consuming about 49% of CPU (Supplementary Note 3). The high computational demand of TEtranscripts reflects its complex modeling for multimapper handling, while PERREO's streamlined approach achieves comparable biological insights with significantly lower computational cost, a valuable advantage, particularly for short-read experiments where repeat mapping is inherently challenging.

| | FeatureCounts + PERREO DEA | TEtranscripts |
|---|---|---|
| Elapsed time (h:mm:ss) | 3:42:41 | 26:56:07 |
| Threads | 14 | 14 |
| CPU cost (%) | 49 | 87 |
| N (samples) | 115 | 98 |
| Contrasts | GBM vs HC, LGG vs HC and GBM vs LGG | GBM vs HC |

**Table 2.** Elapsed time and CPU usage of featureCounts + DEA PERREO workflow versus TEtranscripts.

#### Supplementary Note 5: DEA results from PERREO analysis

Statistical analysis results were exported in tables that contained log2Fold-Change, p-value, adjusted p-value and other additional statistical parameters. The results for each specific analysis were included in Supplementary Tables 3, 4, 5, 6, 7, 8, and 9, where all the contrasts studied were stored in different sheets in each excel file. Here we provide a summary table outlining the main parameters considered in the analysis.

<table>
<tr><th></th><th>Number of DERs</th><th>Final dataset size</th><th>DEA threshold</th><th>Batch effect reduction</th><th>Duplicates removal</th></tr>
<tr><td>ESCA vs HC (plasma)</td><td>9</td><td>840</td><td>|log2FC| > 1 & FDR < 0.05</td><td>Yes (without indicating batch column)</td><td>Yes</td></tr>
<tr><td>ESCA vs HC (plasma)</td><td>49</td><td>840</td><td>|log2FC| > 1 & FDR < 0.05</td><td>Yes</td><td>Yes</td></tr>
<tr><td>GBM vs HC (tissue: T2T)</td><td>262</td><td>2393</td><td>|log2FC| > 1 & FDR < 0.01</td><td>No</td><td>No</td></tr>
<tr><td>GBM vs LGG (tissue: T2T)</td><td>95</td><td>2393</td><td>|log2FC| > 1 & FDR < 0.01</td><td>No</td><td>No</td></tr>
<tr><td>LGG vs HC (tissue: T2T)</td><td>220</td><td>2393</td><td>|log2FC| > 1 & FDR < 0.01</td><td>No</td><td>No</td></tr>
<tr><td>GBM vs HC (tissue: GRCh38)</td><td>313</td><td>2795</td><td>|log2FC| > 1 & FDR < 0.01</td><td>No</td><td>No</td></tr>
<tr><td>GBM vs LGG (tissue: GRCh38)</td><td>129</td><td>2795</td><td>|log2FC| > 1 & FDR < 0.01</td><td>No</td><td>No</td></tr>
<tr><td>LGG vs HC (tissue: GRCh38)</td><td>248</td><td>2795</td><td>|log2FC| > 1 & FDR < 0.01</td><td>No</td><td>No</td></tr>
<tr><td>GBM vs HC (serum EVs)</td><td>20</td><td>1248</td><td>|log2FC| > 0.8 & FDR < 0.05</td><td>Yes</td><td>No</td></tr>
<tr><td>K562 vs H9 (cell lines)</td><td>122</td><td>533</td><td>|log2FC| > 1 & FDR < 0.05</td><td>Yes</td><td>Not applicable</td></tr>
<tr><td>MCF7 vs H9 (cell lines)</td><td>106</td><td>533</td><td>|log2FC| > 1 & FDR < 0.05</td><td>Yes</td><td>Not applicable</td></tr>
<tr><td>Hct116 vs H9 (cell lines)</td><td>74</td><td>533</td><td>|log2FC| > 1 & FDR < 0.05</td><td>Yes</td><td>Not applicable</td></tr>
<tr><td>HepG2 vs H9 (cell lines)</td><td>109</td><td>533</td><td>|log2FC| > 1 & FDR < 0.05</td><td>Yes</td><td>Not applicable</td></tr>
</table>

| | | | | | |
|---|---|---|---|---|---|
| **MCF7 vs Hct116 (cell lines)** | 64 | 533 | $\lvert \text{log2FC} \rvert > 1$ & FDR < 0.05 | Yes | Not applicable |
| **K562 vs HepG2 (cell lines)** | 105 | 533 | $\lvert \text{log2FC} \rvert > 1$ & FDR < 0.05 | Yes | Not applicable |
| **K562 vs Hct116 (cell lines)** | 104 | 533 | $\lvert \text{log2FC} \rvert > 1$ & FDR < 0.05 | Yes | Not applicable |
| **HepG2 vs Hct116 (cell lines)** | 91 | 533 | $\lvert \text{log2FC} \rvert > 1$ & FDR < 0.05 | Yes | Not applicable |
| **MCF7 vs HepG2 (cell lines)** | 111 | 533 | $\lvert \text{log2FC} \rvert > 1$ & FDR < 0.05 | Yes | Not applicable |
| **MCF7 vs K562 (cell lines)** | 106 | 533 | $\lvert \text{log2FC} \rvert > 1$ & FDR < 0.05 | Yes | Not applicable |
| **Tumor vs NMEC (mice tissue)** | 942 | 2651 | $\lvert \text{log2FC} \rvert > 1$ & FDR < 0.05 | No | No |
| **Postreatment vs Pretreatment (dog tissue)** | 2 | 896 | $\lvert \text{log2FC} \rvert > 1$ & FDR < 0.05 | No | Yes |

**Table 3.** Summary of main results and selected criteria for the analyzed datasets.

**Supplementary Note 6: Looking for further validation with consensus_search.py**

A Python script was developed to identify highly conserved candidate primer regions within transposable element families represented in the Dfam database. For each family, profile hidden Markov model (HMM) emission probabilities were parsed to obtain position-specific nucleotide probabilities (A, C, G, T). These probabilities were normalized and used to compute per-position information content (IC). In parallel, the consensus sequence associated with each Dfam family was retrieved. A sliding-window approach (default window size: 80 nt) was then applied across the IC profile, and the window with the highest mean IC was selected as the most conserved region within the consensus sequence. The workflow automatically retrieves Dfam models via the Dfam REST API by resolving family names to accessions, downloading both HMM and consensus FASTA files, anprocessinges them in batch. For each family, the coordinates of the optimal window, its mean IC value, and the corresponding consensus subsequence are reported, providing candidate conserved regions suitable for primer design.
