## Supplementary Tables Information for "PERREO: An integrated pipeline for repetitive elements analysis enables the repeatome expression profiling in cancer"

**Supplementary Table 1.** Overview of publicly available datasets included in this study.

**Supplementary Table 2.** Samplesheets used for the analysis of each dataset.

**Supplementary Table 3.** Differential expression analysis (DEA) results for dataset GSE174302 using the T2T-CHM13 assembly with the options -batch yes and -batch no.

**Supplementary Table 4.** DEA results for dataset GSE147352 using GRCh38 as reference genome.

**Supplementary Table 5.** DEA results for dataset GSE147352 using T2T-CHM13 as reference genome.

**Supplementary Table 6.** DEA results for dataset GSE228512 using T2T-CHM13 as reference genome.

**Supplementary Table 7.** DEA results for the Singapore Nanopore Expression Project dataset using T2T-CHM13 as reference genome.

**Supplementary Table 8.** DEA results for dataset GSE242689.

**Supplementary Table 9.** DEA results for dataset GSE117387.

**Supplementary Table 10.** Top 50 most important repRNAs according to Random Forest models trained on dataset GSE147352 (using GRCh38 and T2T-CHM13 as reference genomes) and on dataset GSE228512.
